# The genetic architecture and evolution of life history divergence among perennials in the *Mimulus guttatus* species complex

**DOI:** 10.1101/2020.07.28.225896

**Authors:** Jenn M. Coughlan, Maya Wilson Brown, J.H. Willis

## Abstract

Ecological divergence is a main source of trait divergence between closely related species. Despite its importance in generating phenotypic diversity, the genetic architecture of most ecologically relevant traits is poorly understood. Differences in elevation can impose substantial selection for phenotypic divergence of both complex, correlated suites of traits (such as life history), as well as novel adaptations. Here, we use the *Mimulus guttatus* species complex to assess if divergence in elevation is accompanied by trait divergence in a group of closely related perennial species, and determine the genetic architecture of this divergence. We find that divergence in elevation is associated with differences in multivariate quantitative life history traits, as well as a unique trait; the production of rhizomes, which may play an important role in overwintering survival. However, the extent of phenotypic divergence among species depended on ontogeny, suggesting that species also diverged in investment strategies across development. Lastly, we show that the genetic architecture of life history divergence between two species is simple, involving few mid to large effect Quantitative Trait Loci (QTLs), and that the genetic architecture of the ability to produce rhizomes changes through development, which has potential implications for hybrid fitness in the wild.

## Introduction

Ecological divergence is a major source of phenotypic differentiation among closely related species [1]. Phenotypic differentiation can promote species coexistence [2], and contribute to reproductive isolation [3]. Yet, the ease at which ecological divergence evolves depends, in part, on the genetic architecture of ecologically adaptive traits. The number of alleles controlling these traits, the distribution of their effect sizes, and gene action (e.g. dominance or epistasis) can impact how easily evolution can proceed [4]. If species diverge in suites of correlated traits, the extent of pleiotropy or close linkage between loci controlling multiple traits may restrict multivariate trait evolution to certain phenotypic axes [5]. Alternatively, if traits are controlled by many loci, only some of which have correlated effects, then evolution may be more flexible [6,7].

Differences between closely related species in ecologically important traits may also change with ontogeny. This can occur simply because as development proceeds, individuals have increased capacity to acquire and devote resources to particular traits [8], and/or because the fitness benefit of said traits may change across ontogenetic stages [9]. For example, investment in antiherbivore defensive traits frequently changes across development in plants [9,10], which may reflect differential fitness effects of herbivore damage across development [10]. However, few studies have quantified the extent of trait divergence between closely related species throughout ontogeny or tracked the genetic architecture of ecologically relevant traits across development (but see [11]).

Differences in elevation between closely related plant species is often associated with phenotypic divergence [8,9]. Plant populations differ predictably in phenotypes along elevational gradients; both within species [10–15], and between close relatives [16–18]. Differences in water availability, herbivory, climate, growing season, or a combination of factors may shape phenotypic evolution along elevational gradients [11,12,14,19,20]. Adaptations to high elevation environments often involve both changes in complex suites of traits-such as life history divergence, including correlated changes in the timing of reproduction, investment in vegetative biomass, and the timing of senescence [10–12,17,21], as well as unique phenotypes, like shorter stature [15,22], changes in leaf physiology [23], or production of belowground morphological structures [24,25]. In particular, investment in belowground biomass in high elevation perennials may be particularly important for overwinter survival.

The *Mimulus guttatus* species complex (Section *Simiolus*, DC, Phrymaceae) comprises about a dozen closely related species that inhabit a broad altitudinal range across the Pacific Northwest and exhibit substantial variation in life history [26]. Considerable effort has been made to characterize life history in this group, though most studies have focused on variation within or between annual species [11,19,27–31], or between annuals and perennials [5,20,32–41]. Little is known about the variation in ecologically associated trait divergence among perennials in this group, although they account for a sizable fraction of species and inhabit ecologically unique habitat relative to annuals.

Here we characterize the geographic and ecological distributions of five perennial taxa within the *M. guttatus* species complex, assess the extent of variation in life history across development using a common garden experiment, and determine if life history variation within and among perennial taxa is correlated with differences in elevation. We then map the genetic architecture of life history divergence between two perennial species that exhibit large differences in life history. We assess the extent to which loci affect multiple traits, and characterize the genetic architecture of a novel life history trait through development. Our findings contribute to our understanding of ecological and trait divergence between closely related species, and shed light on the genetic architecture of both correlated and novel traits.

## Materials & Methods

### Study species

Although the taxonomy of the *M. guttatus* species complex is debated [42,43], at least five distinct perennial taxa are recognized: *M. tilingii* (*sensu lato*)*, M. decorus, M. corallinus,* and two ecotypes of perennial *M. guttatus;* coastal perennials (e.g. *M. grandis*; [42]) and inland perennials (e.g. *M. guttatus sensu stricto*; [42]). *Mimulus decorus* and *M. tilingii* both comprise multiple genetic lineages [42,44,45], but as these lineages form monophyletic groups, we include each as a single taxa.

Perenniality in the *M. guttatus* species complex is defined by the production of stolons. Stolons are horizontally growing, vegetative stems that develop from axillary meristems, and root at each node. Although vegetative at first, stolons have the ability to become reproductive. In addition to stolons, some perennials produce vegetative stems that dive underground early in development (e.g. rhizomes; [46,47]). Rhizomes, like stolons, are horizontally growing, vegetative stems that originate above ground. However, once underground, their morphology changes; they become high branched, white, and repress leaf growth (Fig. S1, Fig. S7). Rhizomes can eventually re-emerge above ground, wherein, their morphology reverts to that of a typical stolon, including the ability to become reproductive. Rhizomes thus represent a unique axis of investment in vegetative biomass and may play an important role in adaptation to harsh environments, but little is known about their ecological or phylogenetic distribution or their genetic basis.

### Estimating elevational distributions

To categorize the elevational niche, we curated occurrence records from Global Biodiversity Information Facility (GBIF; https://doi.org/10.15468/dl.vkfnqr) and collection records (available: http://mimubase.org/) for each species within their native ranges. Given the taxonomic complexity of this group, including the presence of somewhat cryptic species [42,44,45], we further excluded GBIF records that were obtained via iNaturalist. We compiled: 2,067 *M. guttatus*, 317 *M. tilingii*, 133 *M. grandis*, 80 *M. decorus*, and 12 *M. corallina* geographically unique sites that were collected between 1865 and 2019. These ranges largely agree with previously published ranges [46], although we include more northerly occurrences for *M. decorus* compared to [46].We then used the R *Dismo* package to construct random cross-validated maximum entropy species distribution models for each taxa separately (Fig. S2, Fig. S3; see supplemental methods for details). Finally, we randomly sampled 500 locales from the predicted suitable habitat inferred from our maxent models for each species and extracted elevation data for each locale. We merged these predicted occurrences with actual occurrence records, and performed an ANOVA with elevation as the response variable, and species, the data source (e.g. predicted or herbarium occurrence), and their interaction as fixed effects (Table S5; Fig. S4). We then used a pairwise T-test with Holm correction for multiple testing to determine what species differed in elevational niche. All statistical analyses were carried out in R [48].

### Quantifying life history variation

In order to assess trait divergence among perennials of the *M. guttatus* species complex, we performed two common garden experiments. Seeds were cold stratified on moist Fafard 4P potting soil for one week at 4°C to break dormancy, then transferred to long day conditions in the Duke University greenhouses (18h days, 21°C days/18°C nights). On the day of germination, seedlings were transplanted to individual 4” pots and monitored daily.

#### I. Population survey

To study phenotypic differences among perennials, we performed a common garden survey of 199 plants from 40 populations of five perennial taxa (average of 8 populations per species and two maternal families per population; Table S1). For each maternal family, we grew three replicate individuals; one for each developmental time point surveyed.

We surveyed plants at three developmental time points; three weeks post stratification (‘vegetative’ period), the day of first flower (‘early reproductive’ period), and six weeks post stratification (‘late-reproductive’ period). At each time point, one replicate individual per maternal family was unpotted, and belowground biomass was cleaned. Plants were measured for the number of stolons and rhizomes produced. Both stolons and rhizomes (if present) were also measured for several stem traits (i.e. length, width, leaf length, branches per node, highest node of emergence). On the day of first-flower, we scored the date and node of first-flower, corolla length and width, and the length and width of the first true leaf. As the phenotyping was destructive, we discarded individuals after phenotyping. In the few cases in which not all maternal families produced three replicates, we preferentially measured replicates at later developmental times.

#### II. Analysis of population survey

We aimed to assess whether life history differed among perennials of the *M. guttatus* species complex in a common garden. We used PCA to summarize life history variation across multi-trait space. We restricted the PCA to include the total number of stolons, the highest node of stolon emergence, the number and proportion of stems that were rhizomes, and the length, width, and leaf length of above ground stolons. PC1 explained 49.82% of the variation and was the only significant PC according to a broken stick model. We performed linear mixed models using the *lme4* and *car* packages in R [49,50] with PC1 as the dependent variable, species, time point, and their interaction as fixed effects and population as a random effect. We found a significant species by time point interaction (Table S6), and therefore performed linear mixed models for each time point separately with species as a fixed effect and population as a random effect (see Table S6 for full model details). Significance of fixed effects were assessed using Type III Wald’s *χ*^2^ tests using the *Anova* function in the *car* package in R [49,50].

To determine whether variation in life history among populations was related to a population’s source environment we performed additional linear mixed models with the morphological PC1 as the dependent variable, elevation, species, time point, and all possible interactions as fixed effects. Population was included as a random effect. Again, we find a significant species by time point interaction (Table S7). We thus perform submodels for each developmental time point, with PC1 as the dependent variable, species and elevation as fixed effects, and population as a random effect.

Lastly, we evaluated the extent to which life history traits covary within perennial taxa. Strong associations between the timing of reproduction and investment in vegetative biomass have been previously described in *Mimulus [5,34,35]*. To determine if perennials exhibited an association between flowering time and stolon number, we performed a linear mixed model with the node of first-flower as the dependent variable, and the number of stolons and species as fixed effects, and population as a random effect (Table S8).

### Determining the genetic basis of life history divergence

#### I. F2 grow out

We grew 429 F2s derived from a single self-fertilized F1 between *M. decorus* (IMP) and coastal perennial *M. guttatus* (OPB). These species are two of the most phenotypically divergent perennials in the complex (Fig. 1, Fig. 2, Fig. S4), varying in several perennial traits, including the total number of stolons, the presence of rhizomes, and flowering time (Table S3; Fig. S8). We additionally grew 24 replicates of each inbred parental line and F1s. Due to an asymmetric seed barrier between *M. decorus* and *M. guttatus* [44], *M. guttatus* served as the maternal donor for all F1s.

**Fig. 1:**
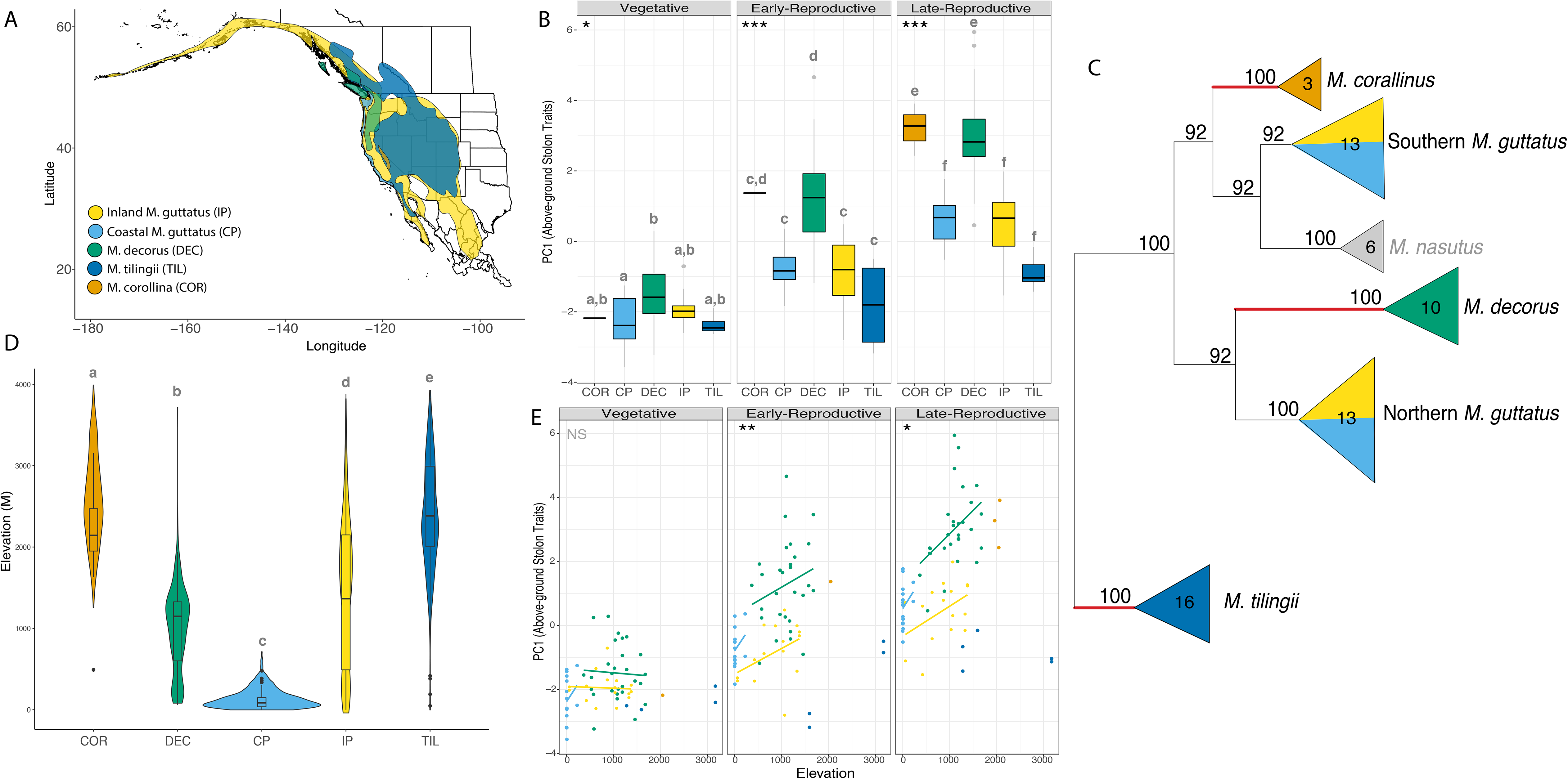
Perennials vary in niche and life history. (A) Geographic ranges of each perennial based on maxent models. (B) Perennials vary in life history, and these differences increase across development (panels). Greater values of PC1 indicate more investment in vegetative biomass. (C) an ML phylogeny of the five perennial taxa (plus annual *M. guttatus* and annual*M. nasutus*) using whole genome sequences. Bolded red branches represent taxa that produce rhizomes at appreciable rates. (D) Differences in elevation niche between perennials. Violin plots represent predicted suitable habitat from maxent models, while interior boxplots are the elevational distribution inferred from herbarium samples. (E) Life history expression within perennial species is correlated with elevation of the source population, but only at later developmental times.

**Fig. 2:**
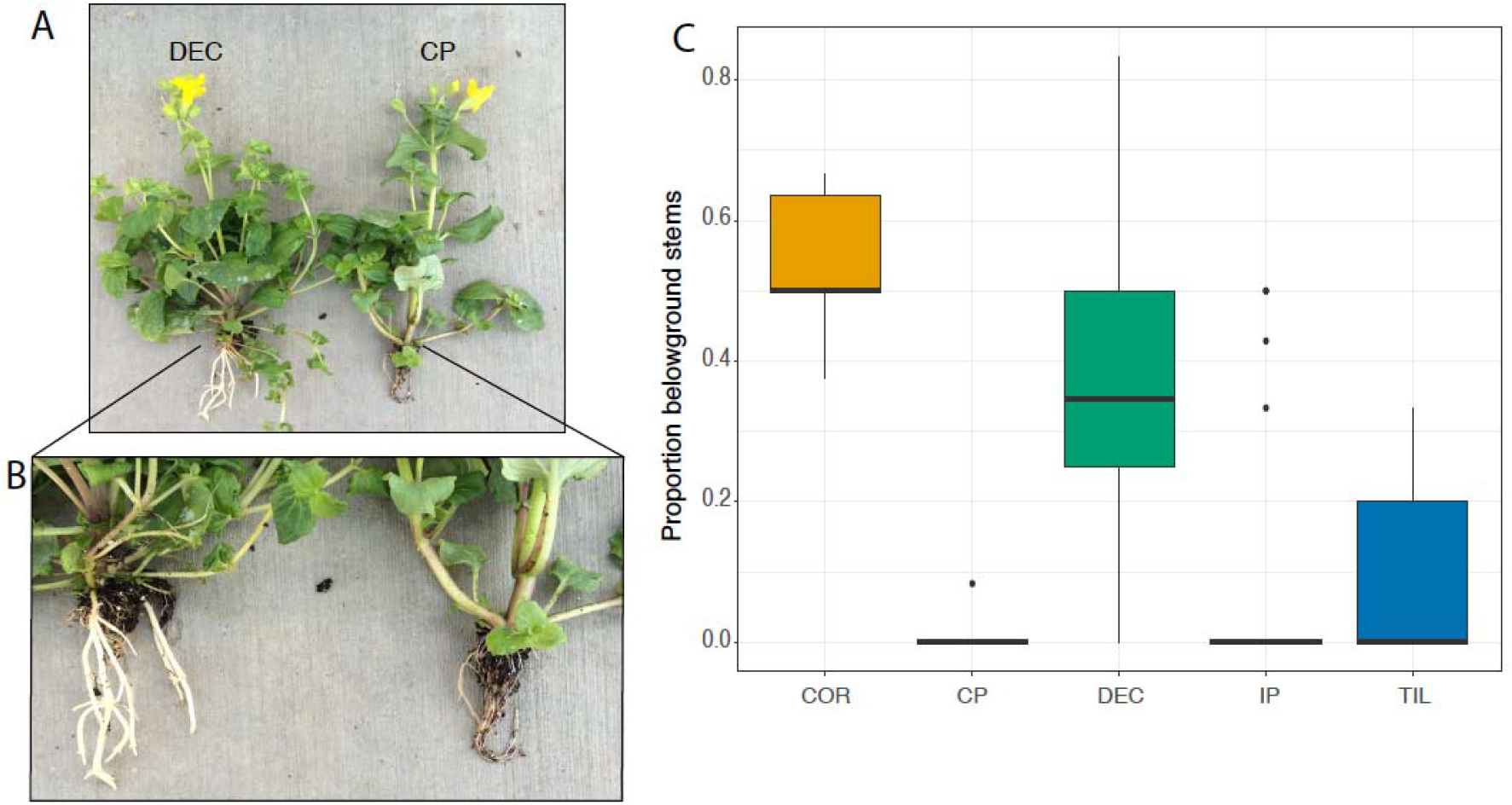
Mid and high elevation perennials make rhizomes. (A) Representative *M. decorus* and coastal perennial *M. guttatus*. Both plants make stolons, but only *M. decorus* produces rhizomes. (B) Close up of the production of stolons in *M. decorus* and a lack thereof in *M. guttatus*. (C) Proportion of rhizomes across development by species. COR= *M. corallinus,* CP= Coastal Perennial *M. guttatus,* DEC= *M. decorus*, IP= Inland Perennial *M. guttatus*, TIL= *M. tilingii.*

Individuals were surveyed at the same developmental time points as above, but were not destructively surveyed. Three and six week post cold stratification individuals were assessed only for the number of stolons and rhizomes. On the day of first flower, individuals were scored for flowering time (date and node of first flower), the length and width of the corolla and first true leaf, the highest node of stolon emergence, and stolons were measured for several traits (length, width, leaf length, number of branches per node, highest node of emergence). A total of 410 F2 individuals were phenotyped, as well as 24 of each parental line and F1. As replicate F1s are largely genetically identical, phenotypic variation among them is primarily due to environmental effects. We thus used the phenotypic variances (σ^2^) within F1s and F2s to estimate broad sense heritability (*h*^*2*^; the proportion of phenotypic variance attributed genetic variance) for each trait as:

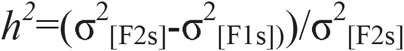

Similarly, we used the phenotypic means (Z) of F1s and parental lines to estimate dominance coefficients for each trait:

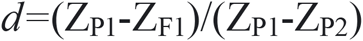

#### II. QTL mapping

We created reduced representation libraries for 384 F2s using a modified Multiplexed Shotgun Genotyping approach with the enzyme *Csp6I* ([52]; see extended methods for details). We aligned and processed reads, then implemented GOOGA to estimate individual genotyping error rates, construct a linkage map, and estimate genotype posterior probabilities for each individual at each marker ([55] see extended methods for details). The final dataset comprised 213 individuals with 1,311 markers. Each individual was genotyped for an average of 974 markers, and 83% of individuals were genotyped at 50% or more markers. Each marker had, on average, genotype information for 74% of individuals.

We used Haley-Knott regressions in the R/*qtl [51,52]* to identify LOD scores, and set significance thresholds using 1,000 permutations. For one QTL on LG4 we find a modest effect on the number of stolons, although this did not pass the genome-wide permutation threshold. We include this QTL in our final map, as single marker analysis using the *fitqtl* function determined there was a significant association (*F*=2.33, *p*=0.033). For traits in which more than one QTL reached significance, we assessed non-additive interactions among QTLs using the *addint* function in R/qtl. Except in one case, we do not recover any significant interactions among QTLs. Lastly, we used these models to calculate the effect size (e.g. *a*, the average difference between homozygotes and heterozygotes), dominance (*d*, the difference between *a* and the mean trait value for heterozygotes), the percentage of phenotypic variance among F2s explained (e.g. %PVE), and the percentage of the parental difference explained by each QTL (e.g. RHE; the relative homozygous effect).

## Results

### Perennial taxa vary in both elevational niche and life history

Perennial taxa of the *M. guttatus* species complex differ in geographic and environmental distributions (Fig. 1, Fig. S2, Fig. S3; Table S5). Four of five taxa are relatively restricted geographically and ecologically, while inland perennial *M. guttatus* is widespread. Of the more restricted taxa, three inhabit almost exclusively mid to high elevation habitats, with *M. tilingii* exhibiting a more broad distribution across the Sierras, Cascades, Rockies, and Pacific Coastal ranges (Fig. S2, Fig. S3) In contrast, *M. corallinus* exclusively inhabits mid to high elevation sites in the Sierras, and coastal perennial *M. guttatus* inhabits low elevation sites along the Pacific coast (Fig. 1; Table S5). Differences in elevation niche were accompanied by differences in climate, with *M. tilingii, M. corallinus* and *M. decorus* inhabiting generally wetter habitats with higher temperature seasonality (Fig. S3; Table S5).

Perennial taxa exhibited substantial variation in life history (as measured by PC1 which explained 49.82% of the variance), and differences between species increased with development (Fig. 1B; Table S6). Greater values of PC1 are associated with greater investment in vegetative biomass via the production of more stolons, stolons that emerge at higher nodes, and stolons that are more branched (Fig. S5). Thus, *M. corallinus* and *M. decorus* tend to make the greatest number of stolons, which emerge at higher nodes and are more branched, while inland perennial *M. guttatus,* coastal perennial *M. guttatus, and M. tilingii* tend to make fewer stolons. A PCA that included reproductive traits (e.g. flowering time, floral size), as well as stolon traits agreed with these results and indicated that perennial taxa that invested more in stolon development also flowered later (Fig. S6).

We aimed to determine if variation in life history in a common garden was related to the elevation of the source population. We find that life history covaried with elevation, but only at later developmental time points (Fig 1; Table S7). Populations originating from higher elevations invested more in vegetative biomass in a common garden; they produced more stolons, and stolons that emerged at higher nodes and are more branched (Fig. 1; Fig. S5). For species that produced rhizomes, higher elevation populations also tended to produce more rhizomes.

We scored the presence of a unique life history trait; the production of rhizomes. Rhizomes are thinner, shorter, and more branched than their above ground counterparts (Fig. S1, Fig. S9). They lack chlorophyll, are substantially more prone to breakage, and repress the growth of vegetative structures relative to stolons (Fig. S1, Fig. S9). Three of five perennial taxa produced rhizomes at appreciable rates (Fig. 2). Although we lack the phylogenetic power to formally test for an association between the presence of rhizomes and habitat, we note that all of the taxa that produce rhizomes are also those that live exclusively in mid to high elevation habitats (Fig. 1, Fig. S7).

### The genetic basis of life history variation among perennials is relatively simple

We constructed an F2 mapping population between *M. decorus* and coastal perennial *M. guttatus* in order to dissect the genetic architecture of life history differences. These two parents vary in several life history traits (Table S3; Fig. S8), including flowering time and size, leaf size, investment in stolons, and the presence of rhizomes, and this is representative of differences between species (Fig. 1, Fig. 2, Fig. S6). The final map consisted of all 14 chromosomes and was 1,232.3cM in length (range: 52.1-125.8cM per linkage group), with an average marker spacing of 1cM.

The genetic architecture of all traits was fairly simple (Fig. 3, Fig. S10). For each of seven traits that differed significantly between parents, we identified one to three significant QTLs with mid to large effects. Individual QTLs explained 9.3% of the trait variance within F2s (range: 4.3-17.2%), and 83.6% of trait difference among parents (range: 7.5−>100%; Table S4) on average. The average cumulative effect of all QTLs across traits was 15.9% of the trait range within F2s, and >100% the trait difference among parents (Table S4). The observation that the cumulative effects of all QTLs for each trait often surpassed the difference between parents is largely due to the fact that several large effect QTLs influenced traits in the opposite direction than predicted from the parental difference (Fig. 3, Table S4), and in line with this observation some F2s exhibited substantial transgressive phenotypes. Of the 12 QTLs we identified, four showed effects opposite to the parental difference, three of which co-localize to LG4.

**Fig. 3:**
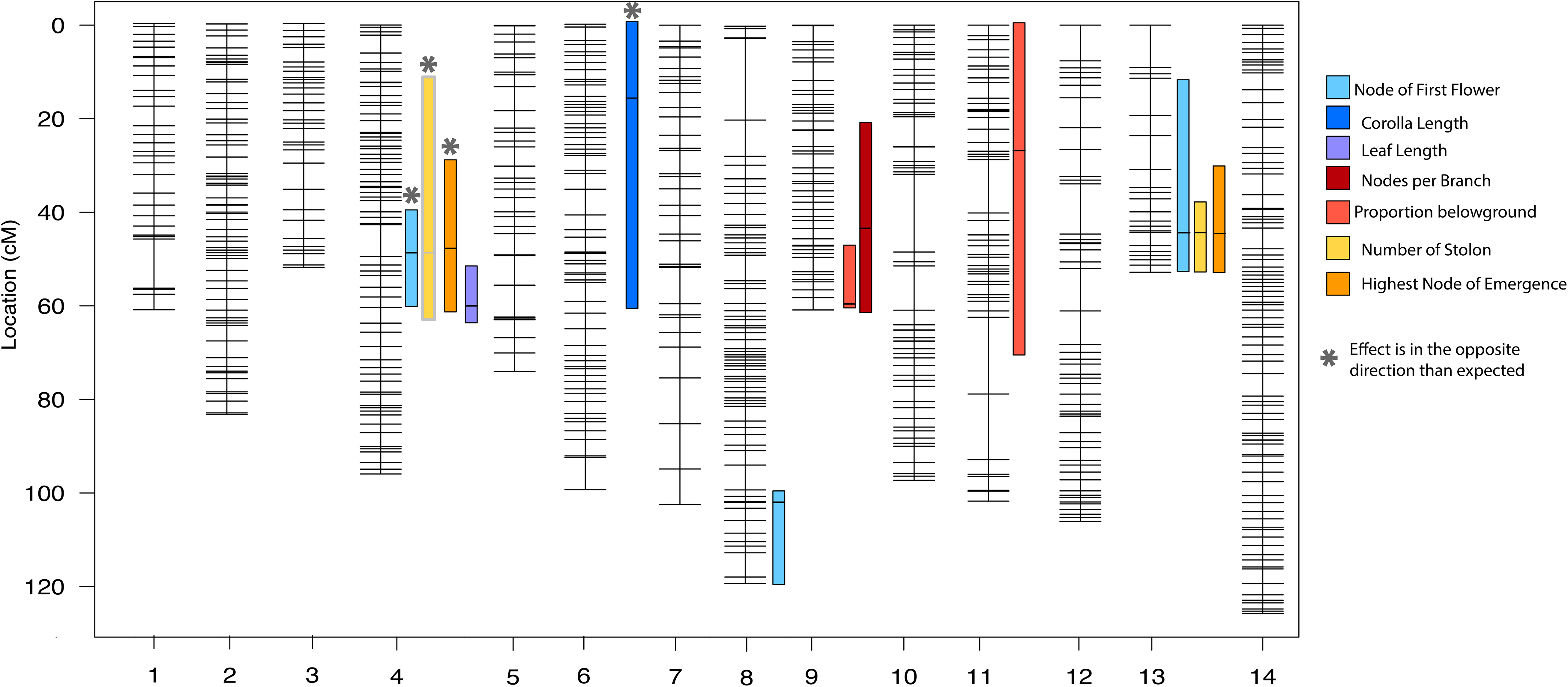
QTL map of life history traits between coastal perennial *M. guttatus* and *M. decorus*. Each colored box represents the 1.5 LOD interval surrounding significant associations (thick horizontal line). Asterisks indicate that the effect of the QTL is opposite of expected. QTL highlighted in grey did not pass a genome-wide significance threshold, but was significantly associated based on individual marker analysis.

We find evidence that two life history traits-flowering time (as measured as the node of first flower) and number of stolons-strongly covary within perennial taxa in the *M. guttatus* species complex (Fig. 4; Table S8). Correlations between these traits persisted among F2s, suggesting that they are genetic (Fig. 4; Table S2). In line with this, QTLs on LG4 and LG13 affected flowering time, the highest node of emergence, and stolon number (Fig. 3; Table S4). The allelic effects of these two QTL are opposite; for LG4, the *M. guttatus* allele is associated with later flowering and more stolons (typically *M. decorus* traits), while the LG13 QTL shows effects in the predicted direction (Fig. 4; Table S4).

**Fig. 4:**
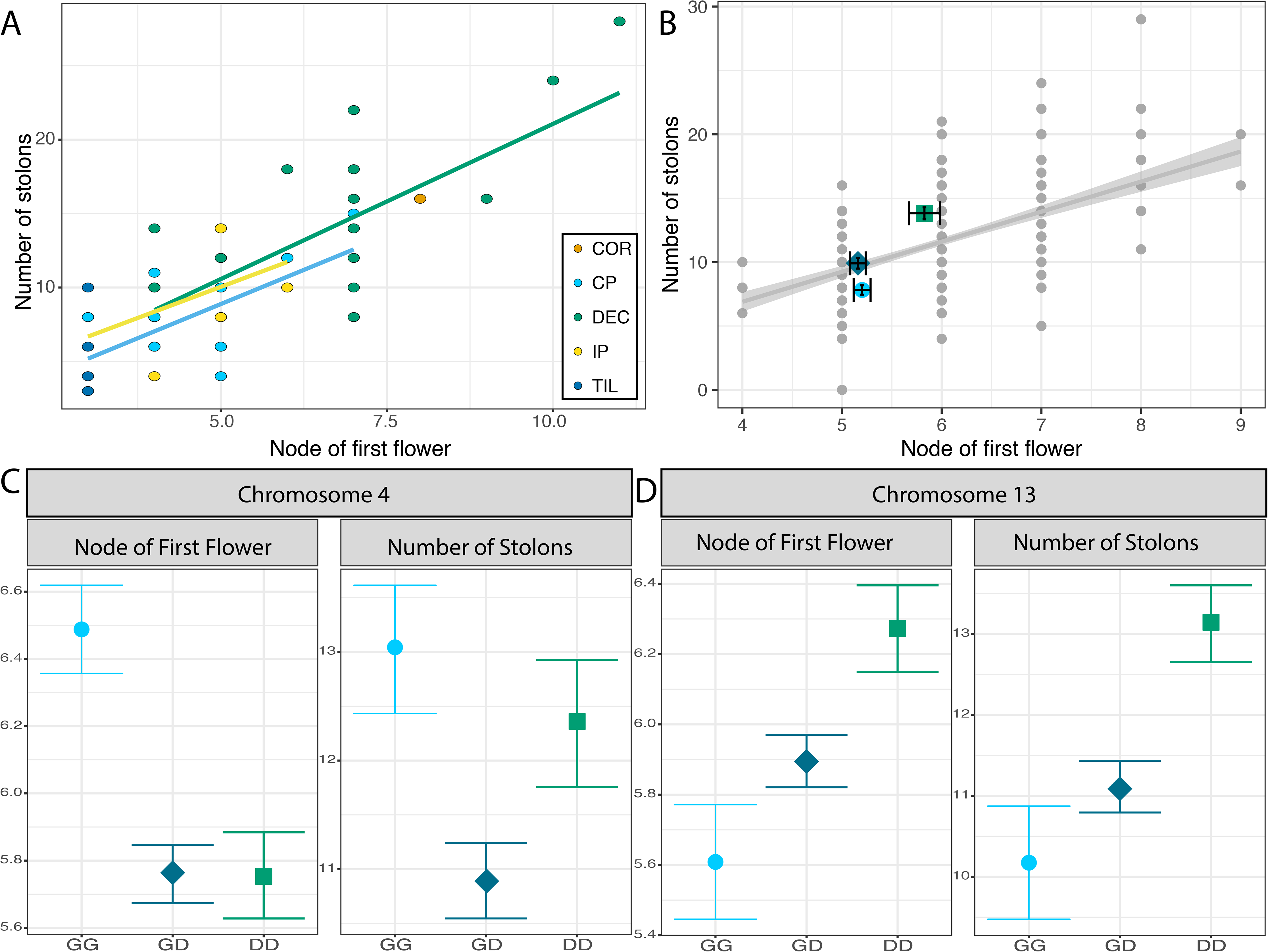
Life history traits covary among perennial species, and share a genetic basis. (A) Phenotypic correlation between flowering time and stolon number across 40 populations of 5 perennial taxa. COR= *M. corallinus,* DEC= *M. decorus,* TIL= *M. tilingii,* IP= inland perennial *M. guttatus,* CP= coastal perennial *M. guttatus.* Lines represent linear regressions for each species. (B) Genetic correlations between flowering time and number of stolons in F2s derived between *M. decorus* and coastal perennial *M. guttatus.* Grey circles= individual F2s, blue circle = the coastal perennial *M. guttatus* parent (OPB), turquoise diamond = F1s, and green square = the *M. decorus* parent (IMP). (C,D) Phenotypic effects of two QTL on flowering time and stolon number. Error bars are standard errors.

Lastly, we find that the production of rhizomes changes through development in hybrids and consequently, so does the genetic architecture. Early on, the ability to produce rhizomes is largely recessive, with few F1s or F2s making rhizomes, and we find no significant QTLs for the ability to produce rhizomes while plants are vegetative (Fig. 5). At reproduction, more hybrids produce rhizomes, but the trait remains recessive. At this point in development, we identify two significant QTLs which exhibit significant non-additive effects (*F*=4.24, *p*=0.0026), such that trait expression in F2s requires the *M. decorus* allele at both loci. However, later in reproduction, most F1s and F2s produce rhizomes. At this point, no interactive effects are detected (*p*=0.89), and a single locus is sufficient to explain ~31% of the parental difference (10.5% of the F2 variation).

**Fig. 5:**
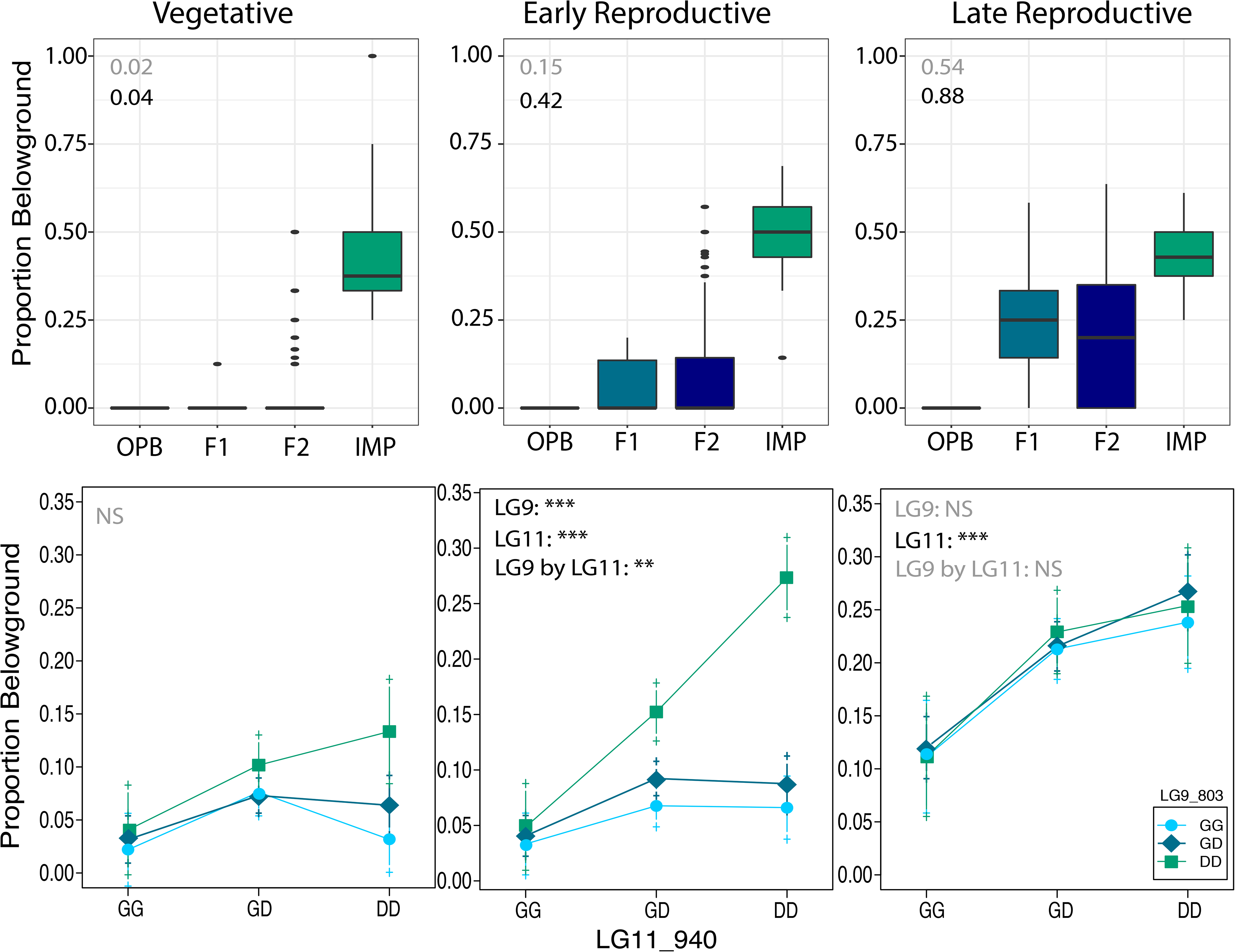
The genetic architecture of rhizomes changes across development. Top row: trait distributions for each parent (OPB= coastal perennial *M. guttatus* and IMP= *M. decorus*), F1s, and F2s. Values in the top left corner indicate the dominance coefficients for the proportion of stems that are rhizomes (top, in grey) and the ability to produce rhizomes (bottom, black). Bottom row: Effect of two markers on the proportion of rhizomes. The significance of each marker and their interaction is listed in the top left corner. G= the coastal perennial *M. guttatus* allele, D= the *M. decorus* allele.

## Discussion

We assessed the extent of divergence in elevational niche and life history among perennial taxa of the *M. guttatus* species complex, and determined the genetic basis of life history differences between two perennial species. We find that perennials exhibited substantial variation in elevational niche and life history, although the extent of phenotypic differences between species depended on ontogeny. Additionally, the presence of a unique life history trait-rhizomes-is largely restricted to species that occur in mid to high elevation habitats, and within species, investment in vegetative biomass is associated with elevation. Together, these suggest that elevation- and covarying factors such as climate, herbivory, or growing season-may partially shape life history evolution within and among perennials of this group. Lastly, we find that the genetic architecture of life history is fairly simple among two closely related perennials. Most traits are largely explained by few mid to large effect QTLs, including the ability to produce rhizomes.

### Elevation may shape life history variation within and among perennials

In a common garden, perennials of the *M. guttatus* species complex exhibit clines in life history along elevational gradients, which is consistent with the hypothesis that elevation- and its associated environmental factors-may, in part, shape life history. Individuals from higher elevations produce more stolons, stolons at higher nodes, and stolons that were more branched. Increased investment in vegetative biomass at higher elevations may increase overwinter survival and promote greater seed production in later years, as each stolon has the potential to become reproductive. In line with this, perennials have higher fitness than annuals in a high elevation common garden [18]. Similarly, adaptive clines in investment in growth and delayed reproduction exist along elevational gradients of annual *M. guttatus* [11,19], as well as other perennial taxa [10,12,13,16,21]. Reciprocal transplants and manipulative common garden experiments will be needed to assess the agents, strength, and targets of natural selection in this system.

*Mimulus tilingii* invested the least in vegetative biomass in our common garden experiment (in agreement with [44,53]), despite inhabiting the highest elevations in nature. *Mimulus tilingii* invests substantially in vegetative biomass in nature [47]. Differences between wild and greenhouse plants likely stem from differences in environment and/or in plant age, wherein high elevation environments may promote investment in more vegetative biomass (as in [15]), and/or naturally collected plants are likely to be older than the plants phenotyped herein and therefore have had more time to invest in vegetative biomass.

We find the taxa that are restricted to mid to high elevation habitats exhibit a novel trait relative to most *M. guttatus* populations; the presence of rhizomes. Rhizomes may afford perennials a greater chance of survival in environments with extreme cold temperatures and sudden freezing, as subsurface soil temperatures tend to be more stable and warmer than ambient air temperatures during freezing events. While uncommon in our experiment, 4/48 inland perennial *M. guttatus* individuals produced rhizomes. These individuals hailed from 2 mid-elevation sites from the Coastal Range and Cascades (1,065m and 866m, respectively). *Mimulus guttatus* can co-occur with *M. decorus* and/or *M. tilingii* at high elevations, and this trait may have introgressed between species. Rhizomes may also have evolved independently in each species or existed as a shared polymorphism in their common ancestor (as discussed in [3,54]). Alternatively, rhizomes may represent an ancestral state, wherein a loss of this character accompanied colonization of low elevation habitats in *M. guttatus*. Given the phylogenetic distribution of this trait across our ML phylogeny, each of these scenarios appears plausible, and further population genomic dissections are required to assess these scenarios.

### The genetic architecture of life history divergence is relatively simple

We find that life history divergence between two perennial species has a relatively simple genetic basis. One to three QTLs explain a significant proportion of the parental difference in most life history traits, and the effects of these QTLs are largely additive. This is simpler than the genetic architecture of life history divergence in other systems [5,28,30,33,55–57]. One explanation for why some genetic architectures are complex versus simple is how natural selection acts on the trait in question [58,59]. Strong, divergent natural selection between populations is expected to produce more simple genetic architectures than stabilizing or balancing selection within populations [58–60]. In the case of life history divergence between coastal perennial *M. guttatus* and *M. decorus*, trait differences may have arisen due to divergent natural selection via elevation (and covarying factors).

We uncover some curious features of the genetic architecture of life history divergence between coastal perennial *M. guttatus* and *M. decorus.* First, we find two QTLs that are associated with three highly correlated traits; stolon number, flowering time, and the highest node of stolon emergence. QTLs that affect both flowering time and stolon number have been described between annual and perennial *M. guttatus [5,33,35]*, and may represent a genetic control of early meristem commitment (e.g. production of floral buds versus stolons; [5,39,61]). The LG4 QTL exhibits opposite allelic effects than predicted by the parental difference. Consequently, some F2s exhibit transgressive phenotypes-delayed flowering and substantially more stolons than either parent. Such extreme phenotypes may have negative fitness consequences in nature, highlighting a potential for reproductive isolation among perennials.

Second, we find that the genetic architecture of the proportion of rhizomes changes with development, in part, because the dominance changes through development. Early in reproduction, the ability to produce rhizomes is recessive, and is controlled by two interacting loci. However, later in development, the ability to produce rhizomes becomes dominant (with the proportion of rhizomes showing minimal dominance), and only one of the previously determined QTLs continues to influence this trait. One potential genetic model for the change in genetic architecture through development is if the LG11 QTL largely controls the ability to produce rhizomes, but timing of its expression is modulated by the LG9 QTL. This has implications not only for understanding the molecular genetic basis of species divergence, but also, potentially for hybrid fitness in nature. While most hybrids eventually produce rhizomes, the fact that the appearance of this trait occurs later in development in hybrids than in *M. decorus* may influence the competitive establishment of rhizomes between hybrids and pure species in high elevation habitats. Further reciprocal transplants will be necessary to assess hybrid vs parental fitness in each species’ habitat.

## Acknowledgements

We thank the Willis and Matute labs, as well as Kate Ostevik and Kieran Samuk for helpful feedback. Maggie Wagner and Aspen Reese provided useful discussions on data analysis. Madison Zamora helped collect plant materials. Dave Lowry, Carrie Wu, and Megan Peterson provided seed stocks. John Kelly and Nic Kooyers gave helpful advice on linkage map construction. This project was funded by an NSF DDIG (DEB-1501758); American Society of Naturalists’ Student research award; and a Society for the Study of Evolution Rosemary Grant Award to JMC, and an NSF Rules of Life award (DEB-1856157) to JW. JMC was funded by an NSF Dimensions of Biodiversity award (DEB-1737752) to Daniel Matute. We are grateful to two anonymous reviews and the editor for helpful feedback that greatly improved the manuscript.

## Author Contribution Statement

JMC conceived of the idea for this project, performed all experiments and data analyses, and wrote the paper. MWB helped with phenotyping and DNA extractions. JW contributed to the intellectual development of this project.

## Data Accessibility Statement

Data for the common garden experiments and sequencing will be made available on Dryad and the NCBI SRA upon acceptance of this manuscript.

## Notes

### Competing Interest Statement

The authors have declared no competing interest.

### Summary of Updates

updated MS draft; re-uploading supplemental tables

## References

1. Endler JA. 1986 Natural Selection in the Wild. Princeton University Press.

2. Schluter D, McPhail JD. 1992 Ecological character displacement and speciation in sticklebacks. Am. Nat. 140, 85–108.

3. Colosimo PF et al. 2005 Widespread parallel evolution in sticklebacks by repeated fixation of ectodysplasin alleles. Science 307, 1928–1933.

4. Orr HA. 2005 The genetic theory of adaptation: A brief history. Nat. Rev. Genet. 6, 119–127.

5. Friedman J, Twyford AD, Willis JH, Blackman BK. 2015 The extent and genetic basis of phenotypic divergence in life history traits in Mimulus guttatus. Mol. Ecol. 24, 111–122.

6. Mitchell-Olds T, Willis JH, Goldstein DB. 2007 Which evolutionary processes influence natural genetic variation for phenotypic traits? Nat. Rev. Genet. 8, 845–856.

7. Wessinger CA, Hileman LC, Rausher MD. 2014 Identification of major quantitative trait loci underlying floral pollination syndrome divergence in Penstemon. Philos. Trans. R. Soc. Lond. B Biol. Sci. 369. (doi:10.1098/rstb.2013.0349)

8. Clausen J. 1949 Genetics of climatic races of Potentilla glandulosa. Hereditas 35, 162–172.

9. Clausen J, Keck DD, Hiesey WM. 1941 Regional Differentiation in Plant Species. Am. Nat. 75, 231–250.

10. Gonzalo-Turpin H, Hazard L. 2009 Local Adaptation Occurs along Altitudinal Gradient despite the Existence of Gene Flow in the Alpine Plant Species Festuca eskia. J. Ecol. 97, 742–751.

11. Kooyers NJ, Greenlee AB, Colicchio JM, Oh M, Blackman BK. 2015 Replicate altitudinal clines reveal that evolutionary flexibility underlies adaptation to drought stress in annual Mimulus guttatus. New Phytol. 206, 152–165.

12. de Villemereuil P, Mouterde M, Gaggiotti OE, Till-Bottraud I. 2018 Patterns of phenotypic plasticity and local adaptation in the wide elevation range of the alpine plant Arabis alpina. J. Ecol. 106, 1952–1971.

13. Buckley J, Widmer A, Mescher MC, De Moraes CM. 2019 Variation in growth and defence traits among plant populations at different elevations: Implications for adaptation to climate change. J. Ecol. 107, 2478–2492.

14. Anderson JT, Wadgymar SM. 2020 Climate change disrupts local adaptation and favours upslope migration. Ecol. Lett. 23, 181–192.

15. Knotek A, Konečná V, Wos G, Požárová D, Šrámková G. 2020 Parallel alpine differentiation in Arabidopsis arenosa. bioRxiv

16. Angert AL, Schemske DW. 2005 The evolution of species’ distributions: Reciprocal transplants across the elevation ranges of Mimulus cardinalis and M. lewish. Evolution 59, 1671–1684.

17. Drummond CS. 2008 Diversification of Lupinus (Leguminosae) in the western New World: derived evolution of perennial life history and colonization of montane habitats. Mol. Phylogenet. Evol. 48, 408–421.

18. Peterson ML, Kay KM, Angert AL. 2016 The scale of local adaptation in Mimulus guttatus: comparing life history races, ecotypes, and populations. New Phytol. 211, 345–356.

19. Kooyers NJ, Colicchio JM, Greenlee AB, Patterson E, Handloser NT, Blackman BK. 2019 Lagging adaptation to climate supersedes local adaptation to herbivory in an annual monkeyflower. Am. Nat. 194, 541–557.

20. Popovic D, Lowry DB. 2020 Contrasting environmental factors drive local adaptation at opposite ends of an environmental gradient in the yellow monkeyflower (Mimulus guttatus). Am. J. Bot. 107, 298–307.

21. von Arx G, Edwards PJ, Dietz H. 2006 Evidence for life history changes in high-altitude populations of three perennial forbs. Ecology 87, 665–674.

22. Boucher FC, Thuiller W, Roquet C, Douzet R, Aubert S, Alvarez N, Lavergne S. 2012 Reconstructing the origins of high-alpine niches and cushion life form in the genus Androsace sl (Primulaceae). Evolution 66, 1255–1268.

23. Ma L, Sun X, Kong X, Galvan JV, Li X, Yang S, Yang Y, Yang Y, Hu X. 2015 Physiological, biochemical and proteomics analysis reveals the adaptation strategies of the alpine plant Potentilla saundersiana at altitude gradient of the Northwestern Tibetan Plateau. J. Proteomics 112, 63–82.

24. Nie X, Yang Y, Yang L, Zhou G. 2016 Above- and Belowground Biomass Allocation in Shrub Biomes across the Northeast Tibetan Plateau. PLoS One 11, e0154251.

25. Chen X-S, Li Y-F, Xie Y-H, Deng Z-M, Li X, Li F, Hou Z-Y. 2015 Trade-off between allocation to reproductive ramets and rhizome buds in Carex brevicuspis populations along a small-scale elevational gradient. Sci. Rep. 5, 12688.

26. Wu CA, Lowry DB, Cooley AM, Wright KM, Lee YW, Willis JH. 2008 Mimulus is an emerging model system for the integration of ecological and genomic studies. Heredity 100, 220–230.

27. Mojica JP, Lee YW, Willis JH, Kelly JK. 2012 Spatially and temporally varying selection on intrapopulation quantitative trait loci for a life history trade-off in Mimulus guttatus. Mol. Ecol. 21, 3718–3728.

28. Troth A, Puzey JR, Kim RS, Willis JH, Kelly JK. 2018 Selective trade-offs maintain alleles underpinning complex trait variation in plants. Science 361, 475–478.

29. Friedman J, Middleton TE, Rubin MJ. 2019 Environmental heterogeneity generates intrapopulation variation in life-history traits in an annual plant. New Phytol. 224, 1171–1183.

30. Fishman L, Kelly AJ, Willis JH. 2002 Minor quantitative trait loci underlie floral traits associated with mating system divergence in Mimulus. Evolution 56, 2138–2155.

31. Ferris KG, Barnett LL, Blackman BK, Willis JH. 2017 The genetic architecture of local adaptation and reproductive isolation in sympatry within the Mimulus guttatus species complex. Mol. Ecol. 26, 208–224.

32. Hall MC, Willis JH. 2006 Divergent selection on flowering time contributes to local adaptation in Mimulus guttatus populations. Evolution 60, 2466–2477.

33. Hall MC, Basten CJ, Willis JH. 2006 Pleiotropic quantitative trait loci contribute to population divergence in traits associated with life-history variation in Mimulus guttatus. Genetics 172, 1829–1844.

34. Hall MC, Lowry DB, Willis JH. 2010 Is local adaptation in Mimulus guttatus caused by trade-offs at individual loci? Mol. Ecol. 19, 2739–2753.

35. Lowry DB, Willis JH. 2010 A widespread chromosomal inversion polymorphism contributes to a major life-history transition, local adaptation, and reproductive isolation. PLoS Biol. 8. (doi:10.1371/journal.pbio.1000500)

36. Twyford AD, Friedman J. 2015 Adaptive divergence in the monkey flower Mimulus guttatus is maintained by a chromosomal inversion. Evolution 69, 1476–1486.

37. Lowry DB, Popovic D, Brennan DJ, Holeski LM. 2019 Mechanisms of a locally adaptive shift in allocation among growth, reproduction, and herbivore resistance in Mimulus guttatus*. Evolution 73, 1168–1181.

38. Lowry DB, Hall MC, Salt DE, Willis JH. 2009 Genetic and physiological basis of adaptive salt tolerance divergence between coastal and inland Mimulus guttatus. New Phytol. 183, 776–788.

39. Baker RL, Diggle PK. 2011 Node-specific branching and heterochronic changes underlie population-level differences in Mimulus guttatus (phrymaceae) shoot architecture. Am. J. Bot. 98, 1924–1934.

40. Baker RL, Hileman LC, Diggle PK. 2012 Patterns of shoot architecture in locally adapted populations are linked to intraspecific differences in gene regulation. New Phytol. 196, 271–281.

41. Rubin MJ, Friedman J. 2018 The role of cold cues at different life stages on germination and flowering phenology. Am. J. Bot. 105, 749–759.

42. Barker WR, Nesom GL, Beardsley PM, Fraga NS. 2012 A taxonomic conspectus of Phrymaceae: a narrowed circumscription for Mimulus, new and resurrected genera, and new names and combinations. Phytoneuron 39, 1–60.

43. Lowry DB et al. 2019 The case for the continued use of the genus name Mimulus for all monkeyflowers. Taxon 68, 617–623.

44. Coughlan JM, Wilson Brown M, Willis JH. 2020 Patterns of Hybrid Seed Inviability in the Mimulus guttatus sp. Complex Reveal a Potential Role of Parental Conflict in Reproductive Isolation. Curr. Biol. 30, 83–93.e5.

45. Sandstedt GD, Wu CA, Sweigart AL. 2020 Evolution of multiple postzygotic barriers between species of the Mimulus tilingii complex. Evolution (doi:10.1111/evo.14105)

46. Nesom GL. 2013 New distribution records for Erythranthe (Phrymaceae). Phytoneuron 67, 1–15.

47. Nesom GL. 2014 Updated classification and hypothetical phylogeny of Erythranthe SECT. Simiola (Phrymaceae). Phytoneuron 81, 1–6.

48. Team RC. 2017 R: A language and environment for statistical computing.

49. Bates D, Maechler M, Bolker B, Walker S, Christensen RHB, Singmann H, Dai B, Scheipl F. 2012 Package ‘lme4’. CRAN. R Foundation for Statistical Computing, Vienna, Austria

50. Fox J, Weisberg S. 2011 An R companion to applied regression. Sage. Thousand Oaks

51. Arends D, Prins P, Jansen RC, Broman KW. 2010 R/qtl: high-throughput multiple QTL mapping. Bioinformatics 26, 2990–2992.

52. Broman KW, Wu H, Sen S, Churchill GA. 2003 R/qtl: QTL mapping in experimental crosses. Bioinformatics 19, 889–890.

53. Coughlan JM, Willis JH. 2019 Dissecting the role of a large chromosomal inversion in life history divergence throughout the Mimulus guttatus species complex. Mol. Ecol. 28, 1343–1357.

54. Pease JB, Haak DC, Hahn MW, Moyle LC. 2016 Phylogenomics Reveals Three Sources of Adaptive Variation during a Rapid Radiation. PLoS Biol. 14, 1–24.

55. Haselhorst MSH, Edwards CE, Rubin MJ, Weinig C. 2011 Genetic architecture of life history traits and environment-specific trade-offs. Mol. Ecol. 20, 4042–4058.

56. Leinonen PH, Remington DL, Leppälä J, Savolainen O. 2013 Genetic basis of local adaptation and flowering time variation in Arabidopsis lyrata. Mol. Ecol. 22, 709–723.

57. Hämälä T, Mattila TM, Leinonen PH, Kuittinen H, Savolainen O. 2017 Role of seed germination in adaptation and reproductive isolation in Arabidopsis lyrata. Mol. Ecol. 26, 3484–3496.

58. Rajon E, Plotkin JB. 2013 The evolution of genetic architectures underlying quantitative traits. Proc. Biol. Sci. 280, 20131552.

59. Remington DL. 2015 Alleles versus mutations: Understanding the evolution of genetic architecture requires a molecular perspective on allelic origins. Evolution 69, 3025–3038.

60. Yeaman S, Whitlock MC. 2011 The genetic architecture of adaptation under migration-selection balance. Evolution 65, 1897–1911.

61. Geber MA. 1990 The Cost of Meristem Limitation in Polygonum arenastrum: Negative Genetic Correlations between Fecundity and Growth. Evolution 44, 799.

