## Supplemental Figs+Files for "The genetic architecture and evolution of life history divergence among perennials in the *Mimulus guttatus* species complex"

**Extended Methods:**

*I. Species Distribution modeling*

To more accurately estimate elevation niche for each perennial taxa in our study, we constructed maximum entropy species distribution models for each taxa separately using the *Dismo* and *raster* package in R (R. J. Hijmans et al. 2017; Robert J. Hijmans and Van Etten 2016). We first narrowed down predictor variables to maximize biological relevance and minimize the covariance among variables. In *Mimulus*, like many plants, seasonal variation in temperature and precipitation likely drive patterns of extinction and colonization and are likely important determinants of range and niche (Sobel 2014) . We therefore include four bioclimatic factors (temperature and precipitation seasonality, mean temperature of the warmest quarter, and mean precipitation of the warmest quarter), as well as elevation in our models. We restricted these models to the spatial extent of each species (plus a 5° buffer) based on occurrence records and previous publications (Nesom 2013).

For each species, we constructed cross-validated maxent models by randomly sampling 2000 pseudo-absence points from the spatial extent of each species. For both pseudo-absence and occurrence data, we downloaded the 4 bioclimatic variables from WORLDCLIM as well as altitude (from the Shuttle Radar Topography Mission) at a minimum resolution of 2.5 minutes of degrees resolution using the *getData* function in the *raster* package in R (Robert J. Hijmans and Van Etten 2016; R. J. Hijmans, Cameron, and Parra 2005). We then randomly assigned 75% of the pseudo-absence and presence data to train the maxent models, reserving 25% of the data to evaluate the performance of each model. For each species, we performed these models 100 times such that both the pseudo-absence points were resampled with each model and a different subset of occurrence records were used to train and test each of the 100 bootstraps. We used AUC to evaluate the models for each species, and report the mean and standard deviation of the 100 bootstrapped models for each species (Fig. S3).

Finally, to assess the predicted elevation niche for each species, we set a threshold for predicted suitable habitat to maximize the sensitivity (e.g. true positive rate) and specificity (e.g. true negative rate) using the *spec_sense* statistic in *Dismo* for each species. For each species, we then randomly sampled 500 geographic locations within the predicted suitable habitat and extracted all 19 bioclim variables from WORLDCLIM as well as altitude at a resolution of 2.5 minutes of degrees. Lastly, to describe predicted elevational differences between species, we combined the predicted and herbarium occurrence records to perform an ANOVA with elevation as the response variable and species, the data source (e.g. predicted or herbarium occurrence), and their interaction as fixed effects. Neither data source, nor the interaction between data source and species were significant for elevation (Table S5), but species did vary significantly in their elevational distribution. To determine what species varied we then performed a pairwise T-test with Holm correction for multiple testing.

*II. Phylogenetic Analyses*

In order to assess the phylogenetic relationship among perennials in the *M. guttatus* species complex, we sequenced three accessions of *M. corallinus*. Briefly, we extracted DNA from bud and leaf tissue using a modified CTAB protocol ((Kelly and Willis 1998); [dx.doi.org/10.17504/protocols.io.bgv6jw9e](https://dx.doi.org/10.17504/protocols.io.bgv6jw9e)). Libraries were constructed by the University of North Carolina Chapel Hill sequencing facility using KAPA-Mantis kits (https://sequencing.roche.com/), and run on an Illumina NovaSeq 6000 platform with 150bp paired-end reads at the UNC sequencing facility. We combined these three accessions with 58 previously published whole genomes from other species within the *M. guttatus* complex (26 *M. guttatus*, 16 *M. tilingii,* 10 *M. decorus*, and 6 *M. nasutus*) and one accession of *M. dentilobus* which was used as an outgroup. For each genome, we trimmed adapters using Trimgalore! (https://github.com/FelixKrueger/TrimGalore), then aligned reads to the *M. guttatus* V2.0 reference genome (https://phytozome.jgi.doe.gov/) using BWA *mem* (Li and Durbin 2009). We sorted and cleaned reads, then marked duplicates using Picard tools (<http://broadinstitute.github.io/picard/>), before genotypes were called individually using *HaplotypeCaller* in GATK (McKenna et al. 2010). Finally, we used *GenotypeGVCFs* to jointly call genotypes for all individuals. The resultant VCF was filtered such that indels were removed, and only bi-allelic sites that had a minimum coverage of 5x, a minimum genotype quality (GQ) of 30, a minimum quality of (minQ) of 30 and at least 47 of 62 (75.8%) of individuals were genotyped were retained using VCFTools (Danecek et al. 2011). We then inferred a maximum likelihood phylogeny using IQtree V2.0.3 (Nguyen et al. 2015). under a TVM+F+R4 model, which IQtree identified as the best fit model using BIC. In total, this ML tree was built using 172,039 sites, 37,294 were parsimony informative. We also performed 1,000 ultra-fast bootstraps in IQtree to assess branch support.

*III. QTL mapping: Library construction, sequencing, read processing, linkage map construction, genotype estimation*

We extracted DNA from F2 individuals using a modified CTAB protocol ((Kelly and Willis 1998); [dx.doi.org/10.17504/protocols.io.bgv6jw9e](https://dx.doi.org/10.17504/protocols.io.bgv6jw9e)). We created libraries for 384 F2s and each parental line using a modified Multiplexed Shotgun Genotyping approach with the enzyme Csp6I ((Andolfatto et al. 2011); [dx.doi.org/10.17504/protocols.io.bbjbikin](https://dx.doi.org/10.17504/protocols.io.bbjbikin)). All eight libraries were split across 2 lanes of Illumina 4000 High Seq at the Duke Sequencing Facility, for an average of 97.4 million reads per library (~2 million reads per F2 individual).

We demultiplexed reads using Stacks 2.2 (Catchen et al. 2013; Rochette and Catchen 2017). Each file was aligned to the V2.0 *M.guttatus* reference genome (https://phytozome.jgi.doe.gov/) using BWA (Li and Durbin 2009), then cleaned and sorted using Picard tools (<http://broadinstitute.github.io/picard/>). We used GATK to call SNPs for each sample individually, then jointly genotype all individuals (McKenna et al. 2010). We removed indels and included sites with a minimum genotype quality of 30 using VCFtools (Danecek et al. 2011). We polarized SNPs in the F2s based on sites that were alternative homozygotes in the parental inbred lines, and determined genotype markers by using the number of counts for each parental allele across 100kb windows. A window was determined to be homozygous if >95% of reads were of one parent, and heterozygous if the percent of reads were between 0.1 and 0.9. We filtered the dataset to remove individuals with very low genotyping (<50 markers across the genome) and markers with genotype information for <10% of individuals.

We implemented GOOGA (Genome Order Optimization by Genetic Algorithm; (Flagel et al. 2019)) to estimate individual genotyping error rates, construct a linkage map, and estimate genotype posterior probabilities for each individual at each marker (Flagel et al. 2019). GOOGA pairs a Hidden Markov Model with a genetic algorithm to first estimate individual genotype error rates, then order markers within and among scaffolds, and finally assign genotype probabilities, given both the individual genotype error rates and likelihood of recombination between markers. First, we used genotypes based on individual SNPs to assign preliminary genotypes in 100KB windows for each individual. A preliminary windowed genotype was only called if that genotype window had at least 1 SNP and at least 5 reads. Windows were deemed homozygous if they contained 90% calls for a single parent, and otherwise were assigned a preliminary status as heterozygous. We then used GOOGA to estimate individual genotype error rates as: (a) the probability that a homozygote is erroneously called a heterozygote, (b) the probability that a homozygote is erroneously called the alternative homozygote, and (c) the probability that a heterozygote is erroneously called a homozygote. In order to construct a linkage map based solely on individuals with high quality genotype calls, we excluded any individual with an error rate over 20% for each of the error types (as in (Flagel et al. 2019)), which resulted in 132 individuals being used to construct the linkage map. We then used GOOGA to calculate recombination rates among markers in order to identify misplaced markers (e.g. markers that are unlinked to those proximate to it) using only the high quality genotyped individuals. This process was run iteratively until all included markers gave reasonable recombination rates (e.g. exhibited some level of linkage with proximate markers). Finally, we used GOOGA to estimate final genotype posterior probabilities, given both the genotyping error rates and the recombination rates defined by the linkage map for all individuals (including those with higher genotyping error rates). We defined final genotypes as those that yielded a posterior probability >/=95%. To filter our final genotype table, we removed individuals with substantial missing data (>90% missing data) or severely distorted overall genotypic compositions (<20% or >80% heterozygous calls). The remaining dataset comprised 213 individuals with 1,311 markers. In total, each individual was genotyped for an average of 974/1,311 markers, and 83% of individuals were genotyped at 50% or more of markers. Additionally, each marker had, on average, genotype information for 74% (median of 75%) of individuals.

**Supplemental Figures**

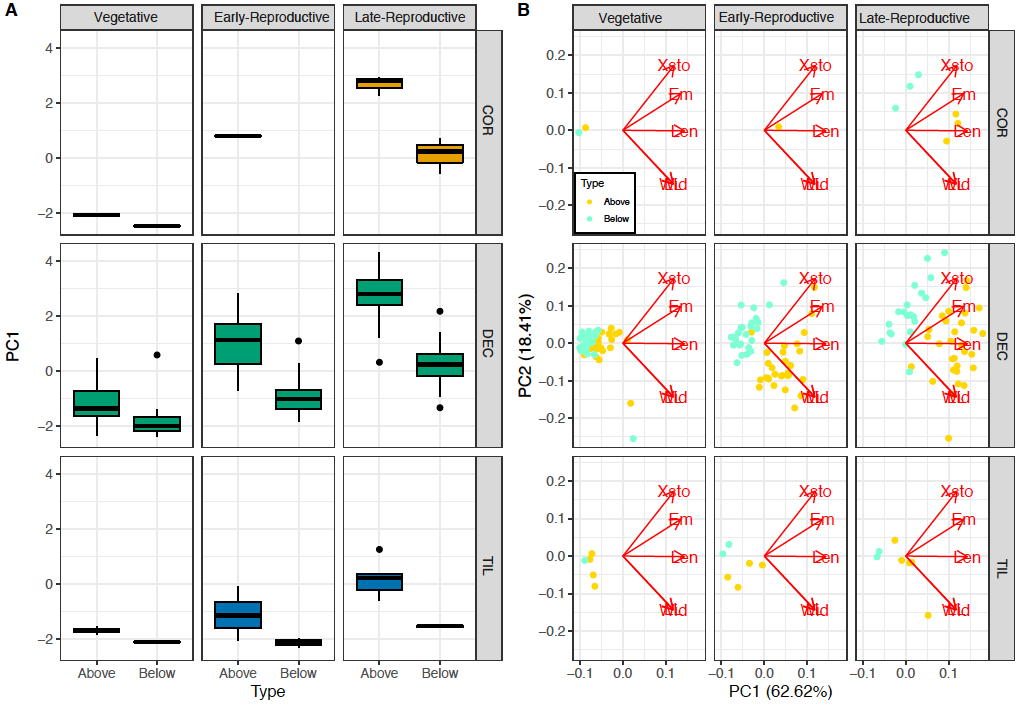

**Figure S1:** Stolons and rhizomes differ in several morphological traits, and these differences grow through development. Analyses were restricted to the three species that made both stolons and rhizomes (*M. decorus, M. tilingii,* and *M. corallinus*). We performed a PCA of all stem traits measured for both stolons and rhizomes (i.e. the highest node of emergence, average branches per node, and the length, width and leaf length of the longest stem). PC1 explained a significant proportion of the variation (62.62%). We find a significant species by time point (*𝜒^2^*=16.627, *df*=2, *p*=0.0002) and marginally significant type by time point interaction (*𝜒^2^*=3.109, *df*=1, *p*=0.077), suggesting that the difference between species and between stolons and rhizomes changed across development. Early in development, stolons and rhizomes are similar and also similar across species (no significant effect of species (*p*=0.17) or type (*p*=0.65)), but differences- both between species and between types of stems- grow as plants develop (early reproductive: significant species effect: *𝜒^2^*=18.3, *df*=2, *p*=0.0001; significant type effect: *χ^2^*=5.695, *df*=1, *p*=0.017; late reproductive: significant species effect: *𝜒χ^2^*=25.079, *df*=2, *p*<0.0001; significant type effect: *𝜒χ^2^*=31.007, *df*=1, *p*<0.0001). For all species, rhizomes tend to be thinner, shorter, more branched, emerge at lower nodes, and have small, embryonic leaves. Qualitatively, they are also achlorotic (i.e. white), and far more prone to breakage than aboveground stolons. (A) Differences among species across development for the first PC of stem morphology. (B) A biplot of stolon morphological differences between species across development, with the loading of each variable denoted by a red arrow. Traits: Xsto= Number of stolons, Em= highest node of stolon emergence, Len= length, Wid= width, LL= stolon leaf length. COR= *M. corallinus,* DEC= *M. decorus,* TIL=*M. tilingii sensu lato.*

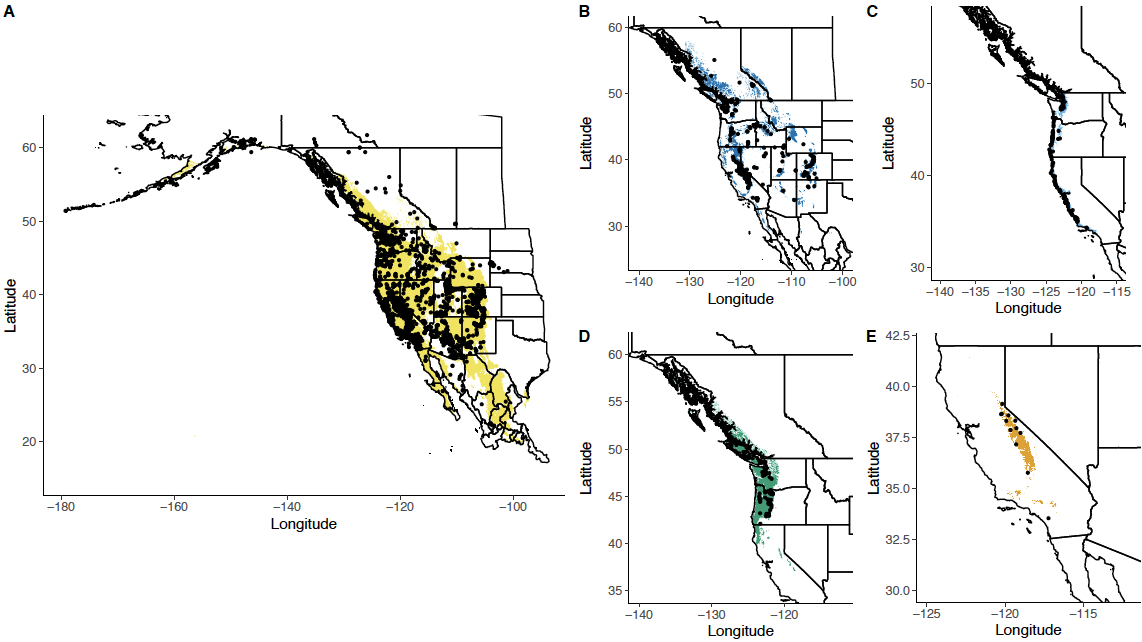

**Figure S2:** Unique occurrence points from Global Biodiversity Information Facility and previous collections (black points) and predicted suitable habitat, as determined by maxent species distribution models for five perennial *Mimulus* taxa: inland *M. guttatus* (A; *M. guttatus sensu stricto,* according to the taxonomic revision prevented in Baker *et al.* 2012), *M. tilingii sensu lato* (B), coastal perennial *M. guttatus* (C; *M. grandis,* according to the taxonomic revision prevented in Baker *et al.* 2012), *M. decorus* (D), and *M. corallinus* (E).

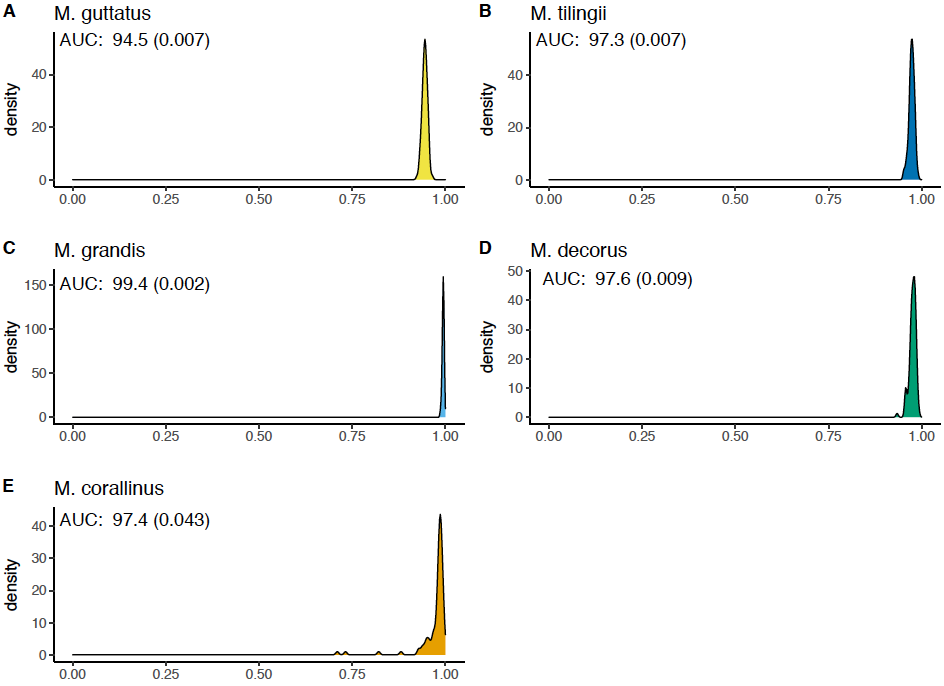

**Figure S3:** Model evaluations for maxent models for each perennial *Mimulus* taxa. Maxent models were evaluated using AUCs (Area Under the Curves), and the distribution of AUCs was estimated by performing 100 bootstrapped models and evaluations. The mean AUC from these bootstraps is given in the top left corner for each taxa, along with the standard deviation in parentheses.

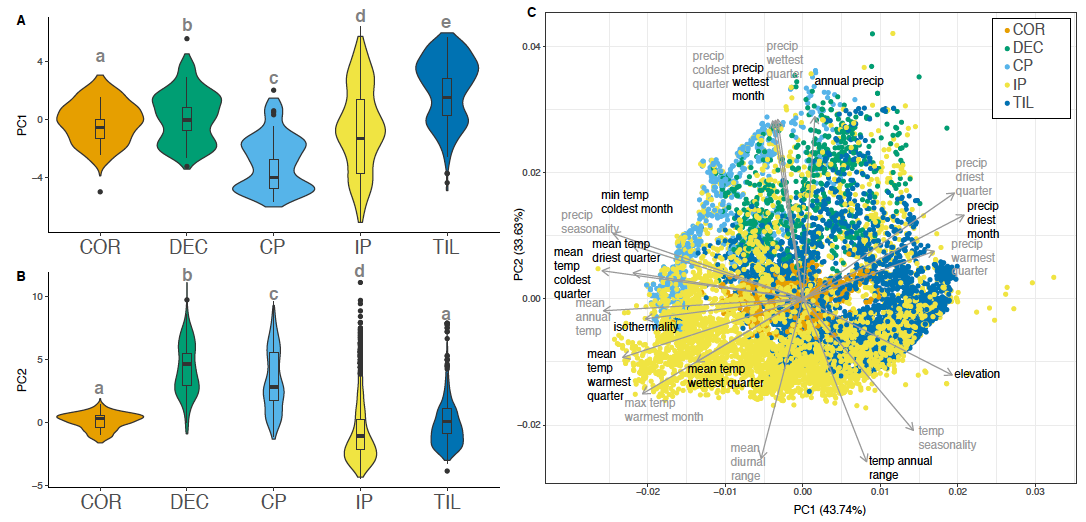

**Figure S4:** Predicted climate envelopes for five perennial *Mimulus* species within the *M. guttatus* species complex. Distribution of PC1 and 2 by species (A,B). For each PC, the distribution of values predicted by maxent models are shown in the violin plot, while the distribution of values for the real occurrence data are shown in the interior boxplot. (C) Biplot of a PCA based on all 19 bioclim variables plus altitude for real occurrence and predicted occurrence data for five perennial species within the *M. guttatus* species complex. PC1 and 2 describe an overwhelming proportion of the variance (43.75 and 33.63%, respectively) and were the only significant PCs according to a broken stick model. Points colored by species. Bioclim variables are differentially colored simply to increase legibility. COR= *M. corallinus,* DEC= *M. decorus,* CP,=coastal perennial *M. guttatus,* IP=inland perennial *M. guttatus*, TIL=*M. tilingii sensu lato.*

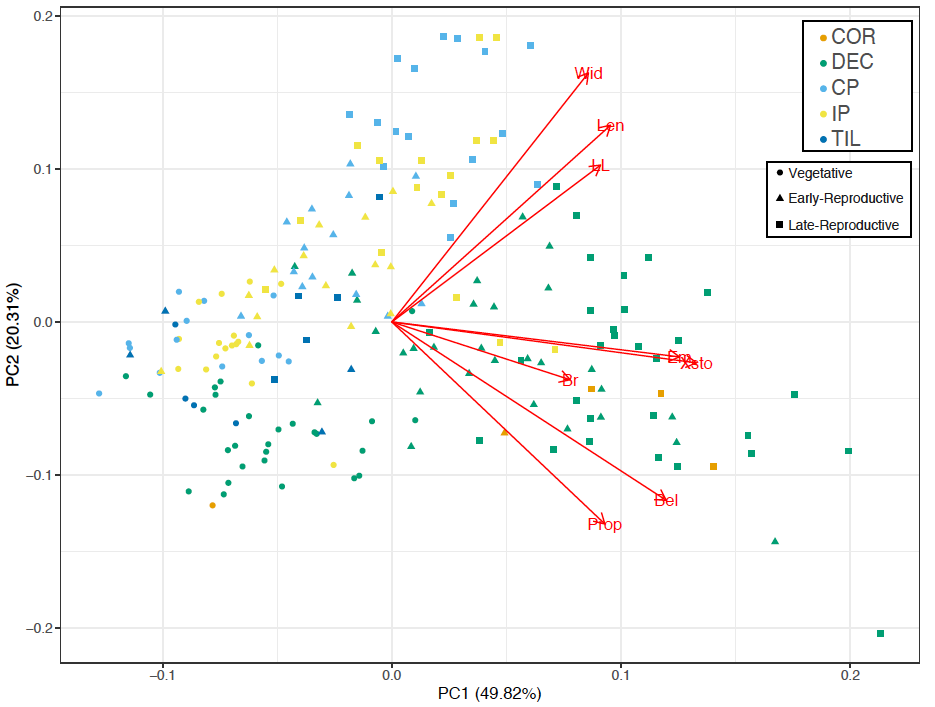

**Figure S5:** Biplot for a PCA involving life history traits across 40 populations of five perennial taxa across three developmental times. LE= stolon length, WI: stolon width, SLL= stolon leaf length, EM= highest node of emergence, ST= the number of stolons, BE= average number of branches per node, and PR= proportion belowground. Colored points refer to the species: COR= *M. corallinus,* DEC= *M. decorus,* CP,=coastal perennial *M. guttatus,* IP=inland perennial *M. guttatus*, TIL=*M. tilingii sensu lato.* Shapes refer to the developmental time point.

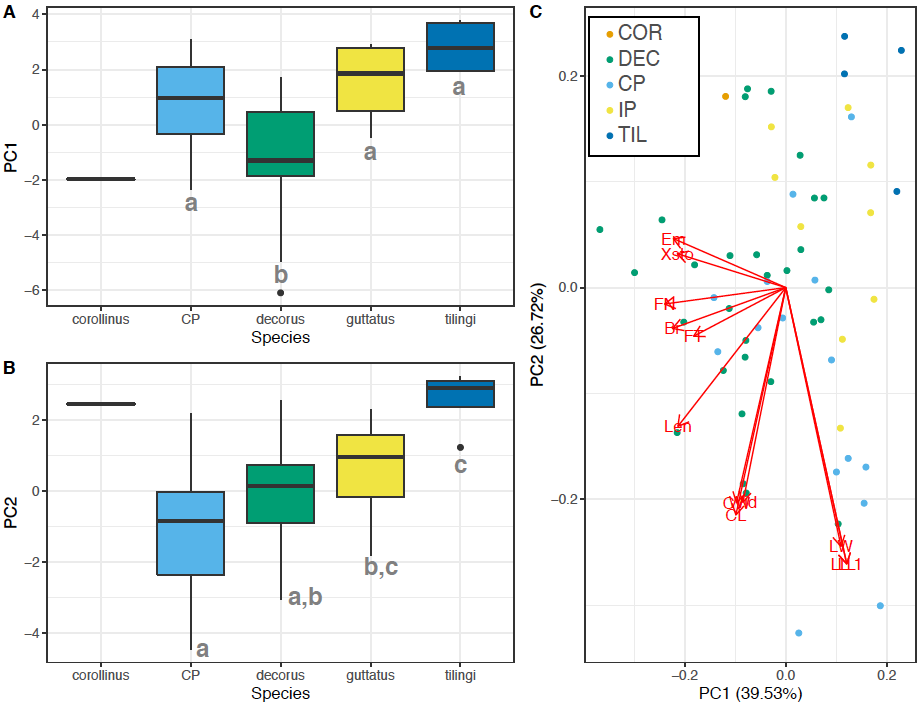

**Figure S6:** An additional PCA of both stolon traits and reproductive traits at only the day of first flower. (A) Values of PC1 across species (39.13% of the variation; Species effect PC1: *𝜒χ^2^*=40.266, *df*=4, *p*<0.0001). (B) Values of PC2 across species (25.77% of the variation; species effect: PC2: *𝜒χ^2^*=18.764, *df*=4, *p*=0.00087). (C) Biplot of the PCA, LE= stolon length, WI: stolon width, SLL= stolon leaf length, EM= highest node of emergence, ST= the number of stolons, BR= average number of branches per node, and PR= proportion belowground, CL= corolla length, CW= corolla width, LL= leaf length, LW= leaf width, FT= days til flower, FN= node of first flower. COR= *M. corallinus,* DEC= *M. decorus,* CP,=coastal perennial *M. guttatus,* IP=inland perennial *M. guttatus*, TIL=*M. tilingii sensu lato.*

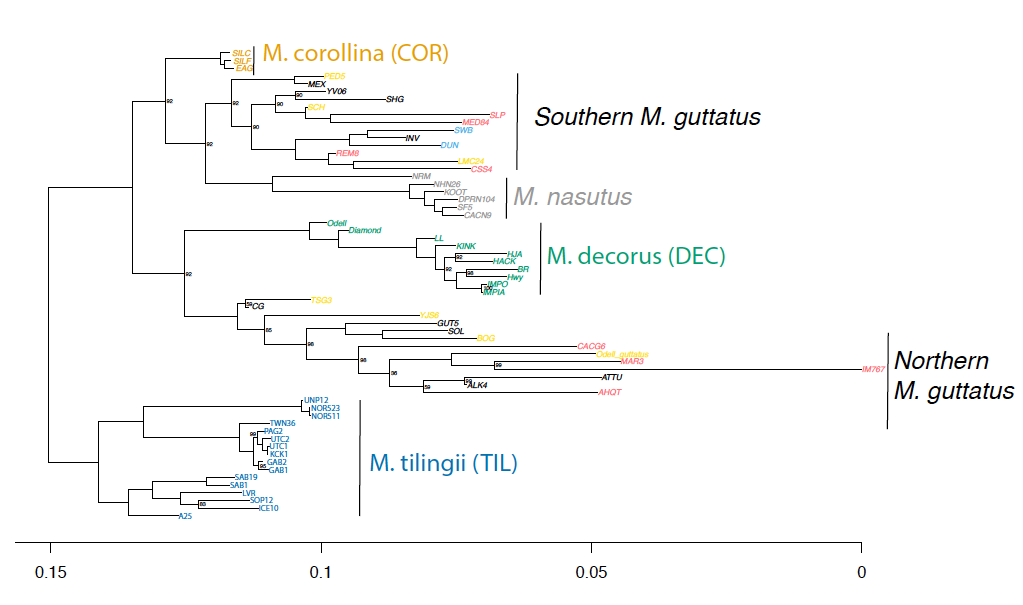

**Fig. S7:** Maximum likelihood phylogeny inferred from IQtree using a TVM+F+R4 model, (which IQtree identified as the best fit model using BIC). Numbers at each node represent the percentage of bootstraps that supported particular relationships, all nodes have 100% support unless otherwise noted. Each species is represented by a unique color, except *M. guttatus* which has been colored by ecotype: coastal perennials (light blue), inland perennials (yellow), annuals (pink), and unknown life history (black).

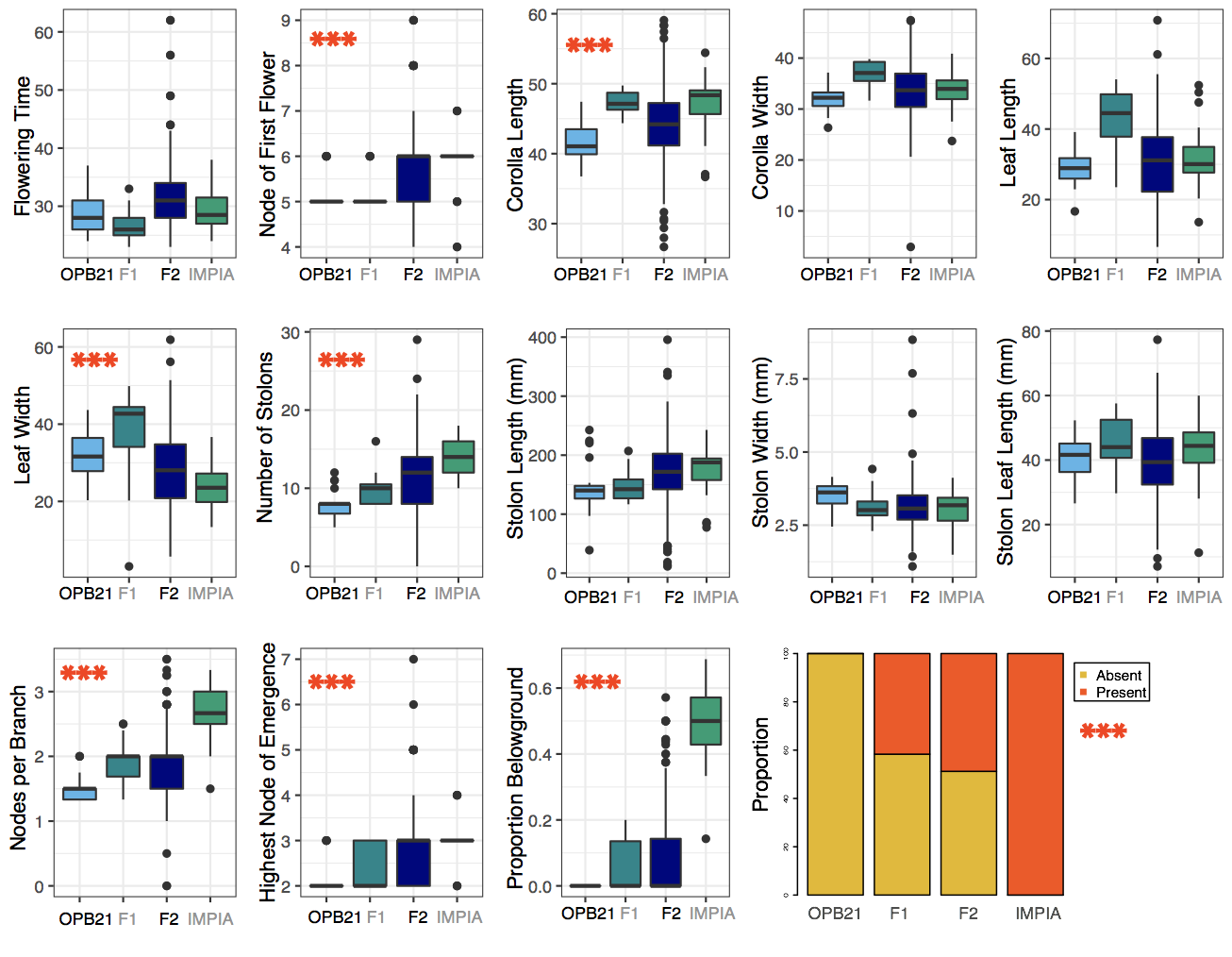

**Figure S8:** Trait distributions for each parental line and their hybrids. Significant differences between parental lines denoted by three red asterisks.

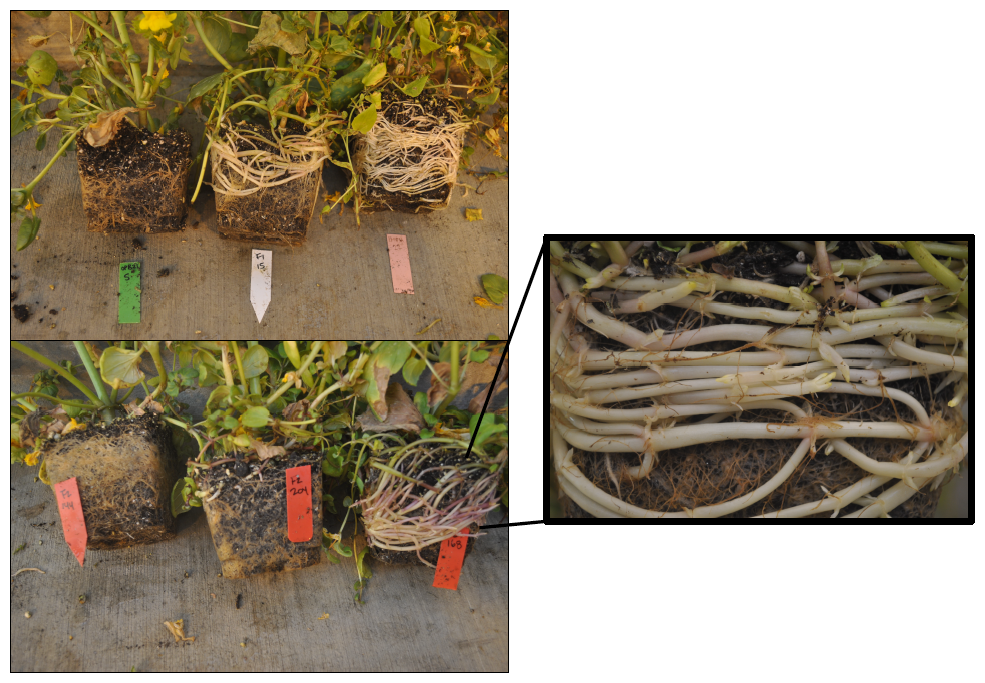

**Figure S9:** Representative pictures of parents and hybrids for the presence of rhizomes very late in development. Top left: Coastal perennial *M. guttatus* (OPB; L), an F1 hybrid (middle), and *M. decorus* (IMP; R). Bottom Left: Three representative F2 individuals. The first produced no rhizomes (L), the second only side branches that delve underground (middle; counted as not producing rhizomes), and the third produced rhizomes (R). Right; close up of rhizomes in a representative F2: note that rhizomes produce embryonic leaves, are white, and branched. Note: photos were taken several weeks after the final phenotyping had occurred.

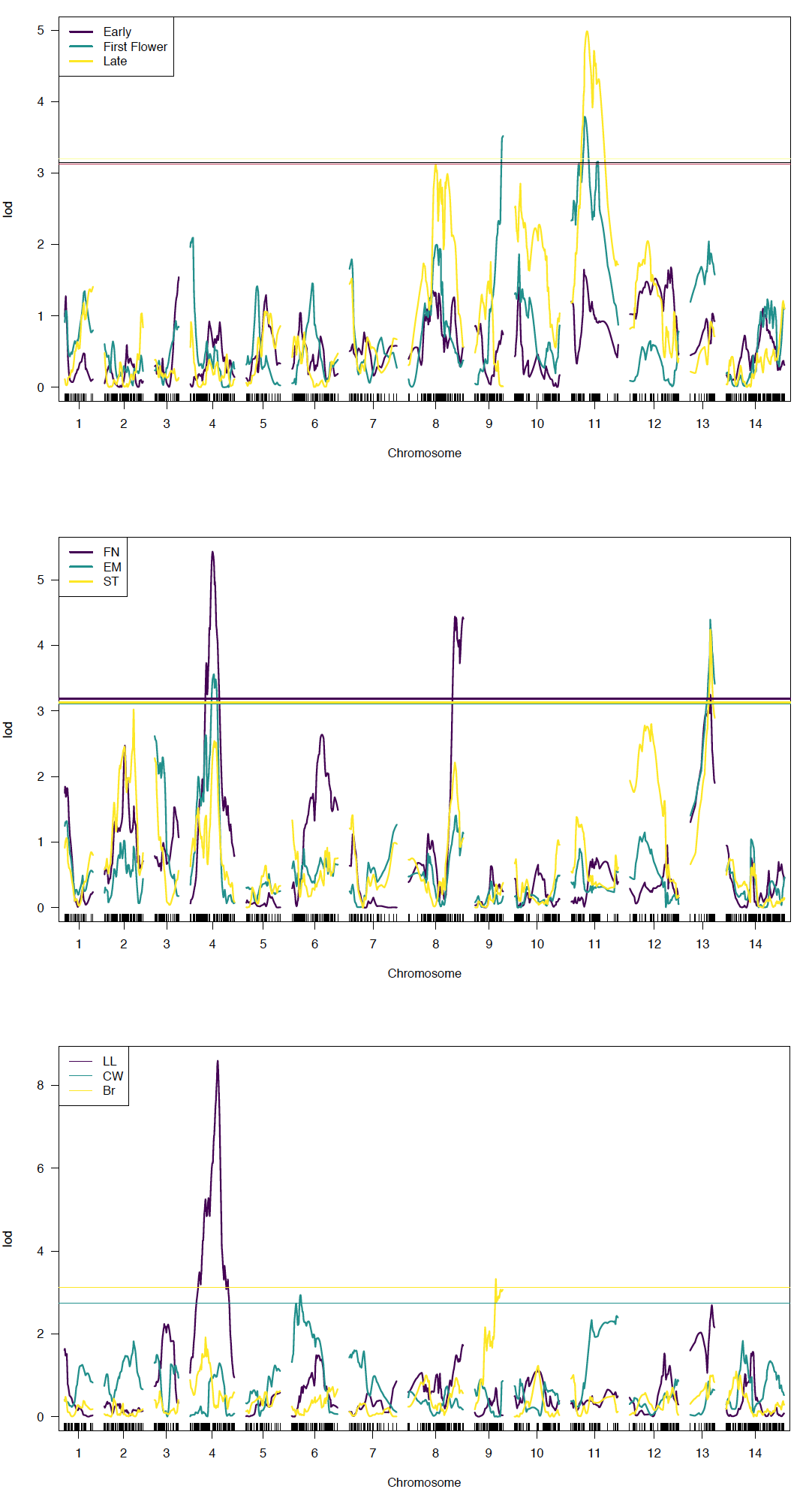

**Figure S10:** Likelihood of Odds (LOD) scores across all chromosomes for all traits that significantly differed between parental lines. Top panel: the proportion of stems that are rhizomes across three developmental time periods: early (3 weeks post stratification), First Flower (the day of first flower), late (6 weeks post stratification). Middle panel: FN (node of first flower), EM (highest node of stolon emergence), ST (number of stolons). Bottom panel: LL (leaf length), CW (corolla width), Br (average branches per node on the longest stolon). Horizontal lines represent that significance threshold as determined by permutation for each trait.

**Table S1:** Seed collections used in a common garden experiment.

| **Population** | **Number of Maternal Families** | **Species** | **Latitude** | **Longitude** | **Elevation (m)** | **Collector** |
| --- | --- | --- | --- | --- | --- | --- |
| EAG | 1 | *M. corallinus* | 38.32 | -119.92 | 2046 | Megan Peterson |
| SILC | 1 | *M. corallinus* | 38.588 | -119.786 | 2066 | Megan Peterson |
| SILF | 1 | *M. corallinus* | 38.664 | -120.219 | 1959 | Megan Peterson |
| 35-48 | 2 | *M.decorus* | 45.305 | -121.667 | 1280 | JMC |
| BM | 2 | *M.decorus* | 44.499 | -121.983 | 1080 | JMC |
| BR | 2 | *M.decorus* | 44.371 | -122.104 | 1189 | JMC |
| DL | 2 | *M.decorus* | 43.158 | -122.133 | 1577 | JMC |
| HACK | 1 | *M.decorus* | 44.40278 | -122.076 | 1189 | JMC |
| HAM | 2 | *M.decorus* | 47.565 | -123.032 | 574 | Carrie Wu |
| HJA | 2 | *M.decorus* | 44.233 | -122.176 | 888 | JMC |
| HWY15 | 2 | *M.decorus* |  |  | 1189 | JMC |
| HWY26 | 2 | *M.decorus* | 45.292 | -121.735 | 1169 | JMC |
| IMP | 2 | *M.decorus* | 44.393 | -122.149 | 1676 | JMC |
| KINK | 1 | *M.decorus* | 44.303 | -121.996 | 1020 | JMC |
| LL | 1 | *M.decorus* | 44.174 | -122.054 | 528 | JMC |
| NS | 2 | *M.decorus* | 44.524 | -121.998 | 966 | JMC |
| ODELL | 2 | *M.decorus* | 43.548 | -121.963 | 1458 | JMC |
| TOLL | 1 | *M.decorus* | 45.321 | -121.907 | 530 | JMC |
| TRILLII | 2 | *M.decorus* | 45.267 | -121.742 | 1102 | JMC |
| WILD | 1 | *M.decorus* | 45.35 | -121.993 | 365 | JMC |
| ZIG | 2 | *M.decorus* | 45.311 | -121.889 | 584 | JMC |
| BCB | 2 | *M.grandis* |  |  | 5 | D. Lowry/K. Wright |
| CVG | 2 | *M.grandis* | 38.372 | -123.055 | 224 | D. Lowry/K. Wright |
| DUN | 1 | *M.grandis* | 43.893 | -124.138 | 0 | D. Lowry/K. Wright |
| HOC | 2 | *M.grandis* | 47.385 | -123.147 | 5 | D. Lowry/K. Wright |
| OPB | 2 | *M.grandis* | 42.464 | -124.423 | 8 | D. Lowry/K. Wright |
| ORO | 2 | *M.grandis* | 35.273 | -120.889 | 5 | D. Lowry/K. Wright |
| OSW | 1 | *M.grandis* | 45.761 | -123.967 | 5 | D. Lowry/K. Wright |
| TSG | 2 | *M.grandis* | 53.419 | -131.916 | 5 | D. Lowry |
| DESCHUTES | 2 | *M.guttatus* | 43.848 | -121.78 | 1377 | JMC |
| DNC | 2 | *M.guttatus* | 41.813 | -124.081 | 33 | Jessica Sely |
| ICE-gutt | 2 | *M.guttatus* | 47.543 | -120.707 | 787 | Jessica Sely |
| NF48 | 2 | *M.guttatus* | 45.202 | -121.615 | 1031 | JMC |
| NTL | 2 | *M.guttatus* | 43.73 | -121.763 | 1328 | JMC |
| SOC | 2 | *M.guttatus* | 41.157 | -122.291 | 626 | Jessica Sely |
| TIH | 2 | *M.guttatus* | 40.876 | -123.756 | 1066 | Jessica Sely |
| WEN | 2 | *M.guttatus* | 47.364 | -120.863 | 866 | Jessica Sely |
| BAG | 2 | *M.tilingii* | 48.873 | -121.688 | 1279 | Carrie Wu |
| SOP | 2 | *M.tilingii* | 38.267 | -119.618 | 3170 | Carrie Wu |
| TWN | 2 | *M.tilingii* | 48.95 | -121.636 | 1594 | Carrie Wu |

**Table S2:** Pearson correlation coefficients between pairs of life history traits across a population survey of 5 perennial taxa (upper right triangle) and within F2s (lower left triangle). FN= Node of first flower; CL= corolla length; LW= leaf width; ST= number of stolons; BR= average branches per node; EM= highest node of stolon emergence; PR= proportion of stems that are rhizomes. Significant correlations (p<0.05) are bolded.

|  | FN | CL | LW | ST | BR | EM | PR |
| --- | --- | --- | --- | --- | --- | --- | --- |
| FN |  | 0.3 | -0.19 | **0.76** | **0.66** | **0.73** | **0.51** |
| CL | 0.09 |  | 0.15 | 0.13 | **0.36** | 0.10 | **0.27** |
| LW | **-0.56** | 0.05 |  | -0.18 | -0.16 | **-0.28** | **-0.31** |
| ST | **0.50** | **0.15** | **-0.28** |  | **0.69** | **0.85** | **0.41** |
| BR | **0.23** | 0.09 | **-0.19** | **0.23** |  | **0.68** | **0.5** |
| EM | **0.65** | 0.11 | **-0.51** | **0.82** | **0.23** |  | **0.37** |
| PR | **0.16** | 0.07 | **-0.16** | **0.29** | 0.06 | **0.21** |  |

**Table S3:** Trait means and standard errors for each parent and F1s, as well as the difference between parents, broad sense heritability (*h^2^*), and dominance of the IMP trait (*d*). Asterisks denote a significant difference between parental means, and was determined by an ANOVA for all traits, except presence/absence, for which we performed a Fisher's Exact Test. Significance thresholds are denoted by asterisks: <0.1 +>0.05<0.05 *>0.01, <0.01 **>0.001, <0.001 ***>0.0001. Traits: days from germination to flower (FT); Node of first flower (FN); Corolla Length (CL); Corolla Width (CW); Leaf Length (LL); Leaf Width (LW); Number of Stolons (ST); Stolon Length (SL); Stolon Width (SW); Stolon Leaf Length (SLL); Average Branches per Node (BR); Highest Node of Stolon Emergence (EM); Proportion of stems that are rhizomes (PR); Presence/Absence of Rhizomes (RH).

| **Trait** | **OPB** | **F1** | **IMP** | **Parental Difference** | **Broad Sense Heritability** | **Dominance (d)** |
| --- | --- | --- | --- | --- | --- | --- |
| FT | 28.88 (0.77) | 26.67 (0.53) | 29.5 (0.74) | 0.62 | 0.75282011 | -3.56 |
| FN | 5.21 (0.08) | 5.17 (0.08) | 5.83 (0.16) | 0.62* | 0.81 | -0.06 |
| CL (mm) | 41.60 (0.51) | 47.38 (0.33) | 47.03 (0.88) | 5.43*** | 0.89 | 1.06 |
| CW (mm) | 32.00 (0.51) | 37.07 (0.26) | 33.49 (0.79) | 1.49 | 0.84 | 3.40 |
| LL (mm) | 28.80 (1.03) | 42.65 (1.83) | 31.84 (1.84) | 3.04 | 0.34 | 4.56 |
| LW (mm) | 32.07 (1.23) | 37.67 (2.24) | 23.74 (1.15) | -8.33* | 0.26 | -0.67 |
| ST | 7.75 (0.37) | 9.83 (0.41) | 13.75 (0.47) | 6*** | 0.71 | 0.35 |
| SL (mm) | 144.05 (8.69) | 147.27 (5.15) | 173.89 (8.88) | 29.84 | 0.75 | 0.11 |
| SW (mm) | 3.52 (0.08) | 3.11 (0.10) | 3.09 (0.12) | -0.43 | 0.52 | 0.95 |
| SLL (mm) | 40.83 (1.36) | 45.26 (1.46) | 43.17 (2.11) | 2.34 | 0.55 | 1.89 |
| BR | 1.50 (0.05) | 1.90 (0.07) | 2.63 (0.09) | 1.13*** | 0.57 | 0.35 |
| EM | 2.17 (0.08) | 2.29 (0.09) | 3.04 (0.11) | 0.87** | 0.68 | 0.14 |
| PR | 0.00 (0) | 0.07 (0.02) | 0.48 (0.25) | 0.48*** | 0.48 | 0.15 |
| RH | 0.00 | 41.67 | 100.00 | 1*** |  | 0.42 |

**Table S4:** Effect size for each QTL for each trait. Significance was determined by 1,000 permutations. CHR and POS are the chromosome and position of the QTLs. LOD is the Likelihood of Odds score. RHE is the percent of the parental difference explained by each QTL (e.g. 'relative homozygous effect'). %PVE is the percent of the F2 variation explained by each QTL. Effect size (a), dominance (d), and PVE were all determined based on the best fit model in *R/qtl*, which for all traits except the proportion of rooting vegetative stems that are rhizomes was purely additive. Expected Direction= Yes/No binary of whether the effect of the QTLs is in the direction expected based on the parental difference. PR= Proportion of rooting vegetative stems that are rhizomes (rather than stolons); FN= Node of first flower; ST= number of Stolons; CL= corolla length; LL= Leaf Length; BR= average branches per node; EM= highest node of stolon emergence. One QTLs is listed in light grey, as this QTLs failed to pass significance thresholds at the genome-wide level, but was significant based on single marker analysis (*F*=2.33, *p*=0.033).

| **Trait** | **CHR** | **POS** | **LOD** | **RHE** | **%PVE** | **a (SE)** | **d (SE)** | **ED?** |
| --- | --- | --- | --- | --- | --- | --- | --- | --- |
| PR | 9 | 60.9 | 3.52 | 23.3 | 13.8 | 0.05 (0.12) | -0.02 (0.02) | Y |
| PR | 11 | 28.8 | 3.79 | 15.4 | 14.3 | 0.05 (0.11) | 0.02 (0.02) | Y |
| FN | 4 | 49 | 5.43 | 117.1 | 10.5 | -0.34 (0.79) | -0.37 (0.12) | N |
| FN | 8 | 100.48 | 4.44 | 57.4 | 9.4 | 0.28 (0.1) | -0.38 (0.13) | Y |
| FN | 13 | 42.97 | 3.25 | 106.2 | 4.3 | 0.25 (0.09) | -0.11 (0.11) | Y |
| ST | 4 | 52 | 2.54 | 7.5 | 5.2 | 0.14 (0.34) | -1.8 (0.50) | N |
| ST | 13 | 43 | 4.24 | 49.2 | 8.7 | 1.5 (0.39) | -0.54 (0.49) | Y |
| CW | 6 | 19 | 2.94 | 86.3 | 6.3 | -2.8 (0.77) | 0.80 (1.19) | N |
| LL | 4 | 60 | 8.59 | 397.7 | 17.2 | 5.4 (1.10) | 7.3 (1.66) | Y |
| BR | 9 | 45.7 | 3.33 | 10.0 | 7.0 | 0.07 (0.06) | -0.31 (0.08) | Y |
| EM | 4 | 51 | 3.56 | 51.1 | 6.4 | -0.11 (0.08) | -0.42 (0.11) | N |
| EM | 13 | 43 | 4.39 | 82.5 | 8.1 | 0.34 (0.09) | -0.1 (0.11) | Y |

**Table S5:** Full model results from ANOVAs to determine if elevation and each significant climate PC differed significantly between species for real occurrence and predicted occurrence data.

| Elevation | | | |
| --- | --- | --- | --- |
| **Response** | ***F*** | **DF** | **p** |
| *Species* | 1050.16 | 4 | <0.001 |
| *Data type* | 37.34 | 1 | 0.31 |
| *Species* Data type* | 2.06 | 4 | 0.08 |
| PC1 | | | |
| *Species* | 415.00 | 4 | <0.001 |
| *Data type* | 12.67 | 1 | <0.001 |
| *Species* Data type* | 0.73 | 4 | 0.57 |
| PC2 | | | |
| *Species* | 973.44 | 4 | <0.001 |
| *Data type* | 36.82 | 1 | <0.001 |
| *Species* Data type* | 1.74 | 4 | 0.14 |

**Table S6:** Model details from Type III Wald’s *χ^2^* test of LMERs to determine if species vary in multivariate trait space across development.

| Full Model: | | | |
| --- | --- | --- | --- |
| **Response** | ***𝜒****χ^2^* | **DF** | **p** |
| *Intercept* | 9.56 | 1 | 0.001 |
| *Species* | 1.67 | 4 | 0.8 |
| *Time point* | 23.26 | 1 | <0.001 |
| *Species* Time point* | 46.04 | 4 | <0.001 |
| Vegetative time point submodel: | | | |
| *Intercept* | 8.57 | 1 | 0.003 |
| *Species* | 12.67 | 4 | 0.01 |
| Early reproductive time point submodel: | | | |
| *Intercept* | 1.48 | 1 | 0.22 |
| *Species* | 55.25 | 4 | <0.001 |
| Late reproductive time point submodel: | | | |
| *Intercept* | 30.5 | 1 | <0.001 |
| *Species* | 97.31 | 4 | <0.001 |

**Table S7**: Full model and submodel by time point details from Type III Wald’s *χ^2^* test of LMERs to determine if elevation within and among species affects trait expression. For each submodel, we excluded species by elevation interactions, as in all cases these interactions were not significant, these models had higher AIC scores and models were not significantly improved by the addition of this interaction term.

| Full Model: | | | |
| --- | --- | --- | --- |
| **Response** | ***𝜒****χ^2^* | **DF** | **p** |
| *Intercept* | 0.19 | 1 | 0.66 |
| *Species* | 2.81 | 4 | 0.59 |
| *Time point* | 12.13 | 1 | <0.001 |
| *Elevation* | 0.07 | 1 | 0.79 |
| *Species* Elevation* | 2.6 | 4 | 0.63 |
| *Species*Time point* | 11.67 | 4 | 0.02 |
| *Elevation* Time point* | 0.45 | 1 | 0.5 |
| *Species* Elevation*Time point* | 4.76 | 3 | 0.19 |
| Vegetative time point submodel: | | | |
| *Intercept* | 6.23 | 1 | 0.01 |
| *Species* | 11.35 | 4 | 0.02 |
| *Elevation* | 0.044 | 1 | 0.83 |
| Early reproductive time point submodel: | | | |
| *Intercept* | 0.30 | 1 | 0.582064 |
| *Species* | 70.63 | 4 | <0.001 |
| *Elevation* | 8.32 | 1 | 0.0039 |
| Late reproductive time point submodel: | | | |
| *Intercept* | 3.53 | 1 | 0.06029 |
| *Species* | 101.42 | 4 | <0.001 |
| *Elevation* | 5.07 | 1 | 0.0243 |

**Table S8:** Results from Type III Wald’s *χ^2^* test of a LMER to assess the correlation between flowering time and stolon number within and between each species included in the population survey.

| **Response** | ***𝜒****χ^2^* | **DF** | **p** |
| --- | --- | --- | --- |
| *Intercept* | 50.26 | 1 | <0.001 |
| *Number of Stolons* | 47.45 | 1 | <0.001 |
| *Species* | 9.11 | 4 | 0.058 |

**References**

Andolfatto, Peter, Dan Davison, Deniz Erezyilmaz, Tina T. Hu, Joshua Mast, Tomoko Sunayama-Morita, and David L. Stern. 2011. “Multiplexed Shotgun Genotyping for Rapid and Efficient Genetic Mapping.” *Genome Research* 21 (4): 610–17.

Catchen, Julian, Paul A. Hohenlohe, Susan Bassham, Angel Amores, and William A. Cresko. 2013. “Stacks: An Analysis Tool Set for Population Genomics.” *Molecular Ecology* 22 (11): 3124–40.

Danecek, Petr, Adam Auton, Goncalo Abecasis, Cornelis A. Albers, Eric Banks, Mark A. DePristo, Robert E. Handsaker, et al. 2011. “The Variant Call Format and VCFtools.” *Bioinformatics* 27 (15): 2156–58.

Flagel, Lex E., Benjamin K. Blackman, Lila Fishman, Patrick J. Monnahan, Andrea Sweigart, and John K. Kelly. 2019. “GOOGA: A Platform to Synthesize Mapping Experiments and Identify Genomic Structural Diversity.” *PLoS Computational Biology* 15 (4): e1006949.

Hijmans, R. J., S. E. Cameron, and J. L. Parra. 2005. “Very High Resolution Interpolated Climate Surfaces for Global Land Areas.” *Journal of Applied Meteorology and Climatology*. <https://rmets.onlinelibrary.wiley.com/doi/abs/10.1002/joc.1276>.

Hijmans, R. J., S. Phillips, J. Leathwick, and J. Elith. 2017. “Package Dismo: Species Distribution Modeling.” *R Package Version 0. 9--3 ed2013*.

Hijmans, Robert J., and J. Van Etten. 2016. “Raster: Geographic Data Analysis and Modeling. R Package Version 2.5-8.”

Kelly, Alan J., and John H. Willis. 1998. “Polymorphic Microsatellite Loci in Mimulus Guttatus and Related Species.” *Molecular Ecology* 7 (6): 769–74.

Li, Heng, and Richard Durbin. 2009. “Fast and Accurate Short Read Alignment with Burrows–Wheeler Transform.” *Bioinformatics* 25 (14): 1754–60.

McKenna, Aaron, Matthew Hanna, Eric Banks, Andrey Sivachenko, Kristian Cibulskis, Andrew Kernytsky, Kiran Garimella, et al. 2010. “The Genome Analysis Toolkit: A MapReduce Framework for Analyzing next-Generation DNA Sequencing Data.” *Genome Research* 20 (9): 1297–1303.

Nesom, G. L. 2013. “New Distribution Records for Erythranthe (Phrymaceae).” *Phytoneuron* 67: 1–15.

Nguyen, Lam-Tung, Heiko A. Schmidt, Arndt von Haeseler, and Bui Quang Minh. 2015. “IQ-TREE: A Fast and Effective Stochastic Algorithm for Estimating Maximum-Likelihood Phylogenies.” *Molecular Biology and Evolution* 32 (1): 268–74.

Rochette, Nicolas C., and Julian M. Catchen. 2017. “Deriving Genotypes from RAD-Seq Short-Read Data Using Stacks.” *Nature Protocols* 12 (12): 2640–59.

Sobel, James M. 2014. “Ecogeographic Isolation and Speciation in the Genus Mimulus.” *The American Naturalist* 184 (5): 565–79.
